# Asymmetric DNA targeting by RNA-guided TIGR-Tas systems

**DOI:** 10.64898/2026.08.11.744272

**Authors:** Kaoling Guan, Nathan M. Appleby, Boyu Huang, Dhruv Champaneri, Griffin Shelley, K. M. Mazharul Alam, Piyush K. Jain, David W. Taylor

## Abstract

Tandem interspaced guide RNA (TIGR)-TIGR-associated (Tas) are RNA-guided defense systems, which use a dual-repeat or stem-loop tigRNA to direct a Tas dimer to DNA targets through RNA-DNA heteroduplex formation between both DNA strands. While previous work has shown the basic principles of DNA targeting, how the guide architecture influences target recognition and whether recognition and cleavage are coordinated across the two RNA-DNA heteroduplexes at a target site remain poorly understood. Here, we combine cryo-electron microscopy, biochemistry, and cell-based assays to investigate two TIGR-Tas effectors: the nuclease-lacking *Peromyscus leucopus* TasA (PlTasA), associated with a stem-loop tigRNA, and the nuclease-active *Salicola phage* TasH (SpTasH). Cryo-EM structures of PlTasA binary and ternary complexes reveal a dimeric scaffold similar to SpTasH and TaTasR with a distinct stem-loop tigRNA architecture and additional peripheral structural elements. Binding assays using DNA substrates containing local bubbles across the spacer-matching region show equivalent bubbles produced position-and spacer-dependent effects, indicating that target engagement is asymmetric in both PlTasA and SpTasH. Kinetic and cryo-EM analyses of SpTasH further reveal stepwise heteroduplex formation, with a partially engaged intermediate that undergoes substantial conformational rearrangements of the second protomer, preceding a fully paired state poised for catalytic activation. Cleavage of the two DNA strands occurs through a coordinated, slow process and productive cleavage requires stringent surveillance of both heteroduplexes. Together, these findings define a conserved, ordered mechanism of bipartite target recognition and activation shared by TIGR-Tas effectors, expanding our understanding of the molecular principles underlying programmable DNA targeting by TIGR-Tas systems.

## Introduction

RNA-guided systems provide a powerful tool for sequence-specific manipulation of nucleic acids and have driven significant advances in biotechnology, particularly in genome editing enabled by CRISPR-Cas (Clustered Regularly Interspaced Short Palindromic Repeats and CRISPR-associated proteins) systems^1–3^. Despite extensive exploration, the diversity of RNA-guided mechanisms in nature remains incompletely understood. Recently, structural and computational mining uncovered a previously unrecognized RNA-guided system with distinct architectures and functions: the tandem interspaced guide RNA (TIGR)-associated (Tas) system, or TIGR-Tas^4^.

TIGR-Tas systems consist of two key components: TIGR arrays, and Tas effector proteins. TIGR arrays are transcribed and processed into 36-or 46-nt tigRNAs. In *Thermoproteota archaeon* TasR (TaTasR) is required for the tigRNA maturation but does not catalyze pre-tigRNA cleavage directly^4^. In contrast, *Salicola phage* TasH (SpTasH) protein is sufficient to process pre-tigRNA into mature tigRNA^5^. Tas proteins are significantly smaller than Cas9 (∼300 versus ∼1,300 amino acids), offering delivery advantages for gene editing applications. Tas proteins contain a conserved RNA-binding nucleolar protein (Nop) domain, which binds RNA in C/D box small nucleolar ribonucleoprotein (snoRNP) complexes^6^, and a coiled-coil domain that serves as a dimerization motif. TasR and TasH additionally possess a RuvC or HNH nuclease domain, respectively^4^. Unlike CRISPR guide RNAs (gRNAs), tigRNAs contain two spacers arranged in tandem, enabling a distinctive targeting mechanism in which each spacer base-pairs with a strand of the target DNA. Additionally, TIGR-Tas systems recognize DNA without requiring a protospacer-adjacent motif (PAM)^4^. Biochemical assays demonstrate that TasR and TasH are RNA-guided DNA endonucleases, with cleavage occuring at defined positions relative to each spacer, generating double-strand breaks. Cryo-electron microscopy (cryo-EM) reveals that TaTasR forms a symmetric dimer that coordinates tigRNA and target DNA, explaining the tandem targeting mechanism^4^. TasA variants lack a nuclease domain, implying roles beyond DNA cleavage, such as transcriptional regulation or competition between mobile genetic elements. These mechanistic features differ fundamentally from canonical CRISPR systems, suggesting distinct principles of RNA-guided targeting and expand the potential toolkit for programmable genome engineering^7,8^ .

Recent studies established the overall architecture and cleavage mechanism of SpTasH^5^, but whether the two heteroduplexes assemble in an ordered manner, how their formation is coordinated, and how nuclease-deficient TasA proteins recognize targets remain unknown. Here, we combine biochemistry, kinetics, mismatch profiling, and cryo-EM to unravel the mechanisms of DNA recognition and effector function in PlTasA and SpTasH. We demonstrate that PlTasA and SpTasH engage target DNA through a tigRNA-directed, asymmetric pathway. SpTasH acts as a slow nuclease whose catalytic activity is tightly coupled to the formation of both RNA-DNA heteroduplexes, thereby promoting coordinated strand cleavage and stringent target discrimination. Structural analyses further reveal a stepwise pathway of heteroduplex maturation, in which pairing of Spacer B represents a late-stage event that drives conformational rearrangements in protomer B. Collectively, these findings expand our understanding of programmable DNA targeting by TIGR-Tas systems and establish a mechanistic framework for the rational engineering of RNA-guided DNA-binding and DNA-cleaving tools.

## Results

### Distinct features of DNA recognition by nuclease-lacking PlTasA

Most TIGR-Tas loci identified to date are associated with dual-repeat arrays, with a subset containing stem-loop structures⁴. How a stem-loop guide directs target recognition has remained unexplored. To address this, we determined cryo-EM structures of a nuclease-lacking TasA from *Peromyscus leucopus* (PlTasA), which is associated with a stem-loop array⁴. We determined a 3.06 Å structure of the target-bound ternary complex and a 3.90 Å structure of the PlTasA-tigRNA binary complex. The structures reveal a dimeric scaffold, with each protomer engaging through a coiled-coil domain and flanked by an RNA-binding Nop domain (**Fig. 1a,b**). Consistent with the established nomenclature for SpTasH and TaTasR⁴,⁵, we refer to the 5’ and 3’ spacers of the tigRNA as Spacer A and Spacer B, respectively. Similarly, we designate the protomer contacting the tigRNA 5’ and 3’ ends as protomer A (the other as protomer B), and the strand pairing Spacer A as Strand A. An N-terminal loop is not resolved in either map, likely owing to intrinsic domain flexibility (**Fig. 1a,b**). In the ternary complex, the target DNA is locally unwound across a continuous region, where both Spacer A and Spacer B form a separate RNA-DNA heteroduplex on opposite strands. Each end of this unwound region is structurally equivalent to the R-loop junction described for SpTasH⁵ (**Fig. 1e**).

**Figure 1.**
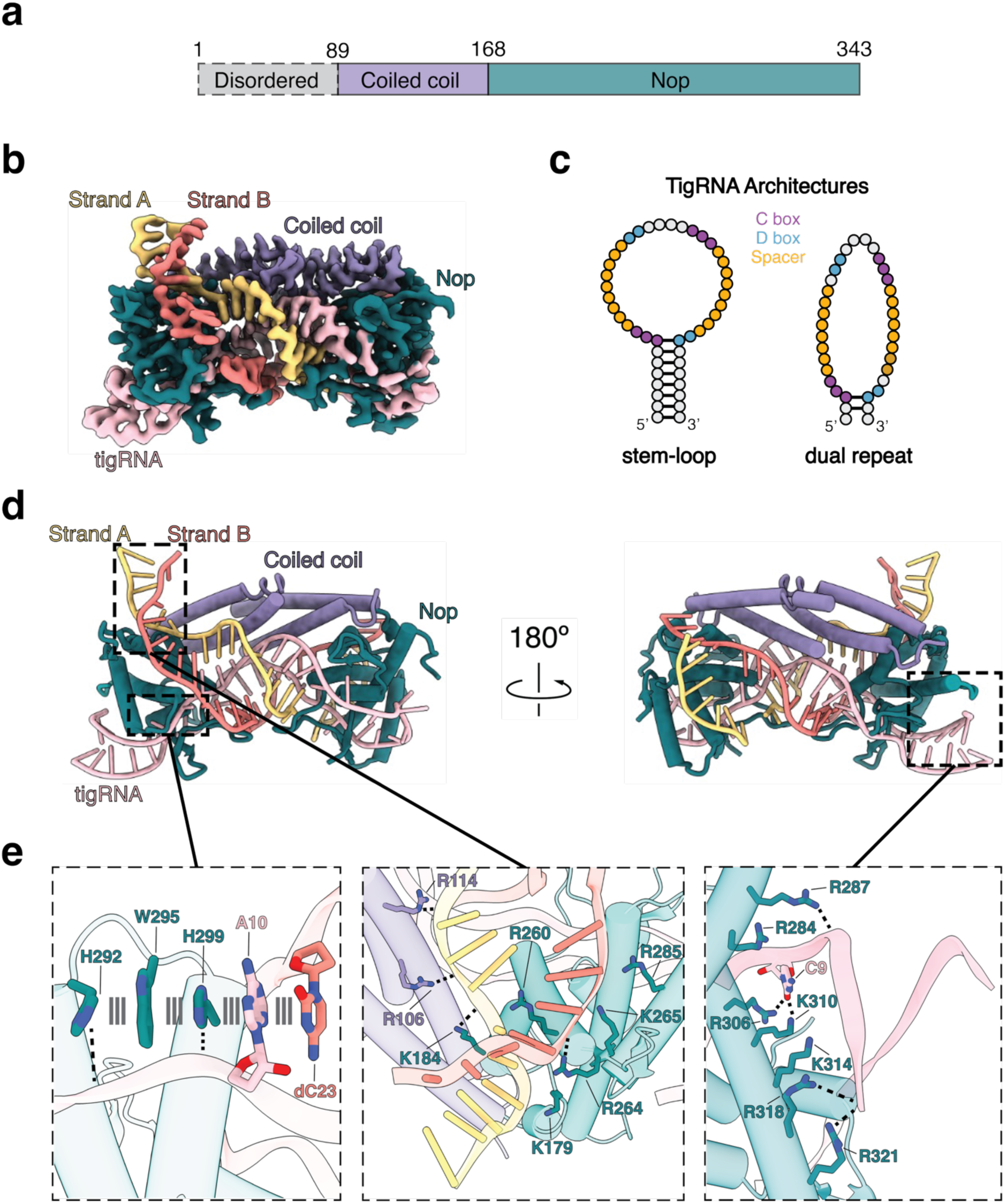
Architecture of the target-bound PlTasA complex. **a.** Domain organization of PlTasA. The unresolved region is colored gray, while the coiled-coil and Nop domains are colored purple and teal, respectively. **b.** EMReady2-processed cryo-EM map of the PlTasA ternary complex. The map is colored based on proximity to the modeled domains shown in **a**. The tigRNA is colored pink, while DNA strands A and B are colored gold and orange, respectively. **c.** Schematic of the two tigRNA architectures, stem-loop and dual-repeat, based on those of PlTasA (left) and SpTasH (right), respectively. Spacers are colored orange; the C and D box motifs are colored purple and light blue, respectively. **d.** Structural model of PlTasA ternary complex colored based on proximity to the modeled domains shown in a. The tigRNA is colored pink, while DNA strand A and B are colored gold and orange, respectively. **e.** Zoomed-in views of three regions. (Left) The extensive stacking interaction between Nop domain residues (H292, W295, H299), the tigRNA (A10), and the displaced DNA strand (dC23). (Center) The unwinding junction of PlTasA, with nearby basic residues positioned to stabilize the unwound DNA. (Right) A cluster of basic residues that interact with the tigRNA stem region. Contacts are designated by dashed lines.

PlTasA shares the overall architecture with the nuclease-active effectors SpTasH and TaTasR. Structural superposition showed that the PlTasA CC-Nop core closely resembles those of TaTasR and SpTasH (**Supplementary Fig. 1**). While the individual CC and Nop domains are structurally conserved across all three effectors, differences in their relative orientations make the overall architecture of PITasA more similar to TaTasR than to SpTasH. Each PlTasA RNA-DNA heteroduplex comprises nine base pairs, matching TaTasR but one base pair longer than SpTasH. Conservation extends beyond the overall architecture to the unwinding junction, where each effector employs a conserved aromatic-basic residue pair on the α4 helix. The aromatic residue stacks against the first tigRNA nucleotide paired with DNA, whereas the basic residue intercalates into the adjacent DNA duplex. In PITasA, these residues correspond to F252 and R260, compared with W203 and K211 in SpTasH, and Y238 and K233 in TaTasR (**Supplementary Fig. 2**). Although the aromatic residue varies between phenylalanine, tryptophan, and tyrosine, mutational analysis of SpTasH showed that substituting W203 with either tyrosine or phenylalanine preserves DNA cleavage⁵, indicating that aromatic stacking rather than residue identity is the defining feature. These conserved structural features suggest that TIGR-Tas effectors use a common mechanism to stabilize the unwinding junction despite differences in nuclease domains and guide architectures.

Beyond the conserved recognition core, PlTasA carries three accessory hairpin elements surrounding the guide-binding region and DNA unwinding junctions (**Supplementary Fig. 2**). Hairpin A (T222-A232) and its following loop (ending at T245) occupies the same position as a loop in TaTasR that caps the displaced strand at each junction, consistent with the overall structural similarity between PlTasA and TaTasR. Unlike the β-hairpin insertion of SpTasH, however, it does not recruit the nuclease domain and instead makes only limited contacts with the RNA-DNA heteroduplex⁵. Hairpin B (L198-R209) lies adjacent to the box D motifs, where R209 contacts a box D nucleobase in one protomer. Hairpin C (I263-A273) projects from helix α4 toward the DNA unwinding junction, positioning R264 to contact the phosphate backbone at the point of DNA strand separation, suggesting a role in stabilizing target engagement (**Fig. 1e**).

Despite its distinct stem-loop tigRNA architecture, PlTasA engages its guide through similar protein contacts to those of the dual-repeat systems. In SpTasH and TaTasR, the 5’ and 3’ ends of the tigRNA form a short, locally base-paired element adjacent to the box C and D motifs, whereas in PlTasA they form a single extended stem (**Fig. 1c**). Recognition of Box C is largely conserved, with R306 and K310 contacting nucleobases directly and R287 engaging the phosphate backbone (**Fig. 1e, Supplementary Fig. 2**). The conserved box C adenine (A10) stacks with H292, W295, H299, and the DNA nucleotide dC23 (**Fig. 1e**). This stacking network is more extensive in PlTasA than in SpTasH or TaTasR. The longer stem is further stabilized by a basic patch comprising R318, R321, and K314, which contacts the phosphate backbone in a region inaccessible to the shorter dual-repeat guides (**Fig. 1e**). Together, these contacts preserve the positioning of box C relative to the DNA unwinding junction as in the dual-repeat systems, while accommodating the extended stem-loop architecture.

Comparison of the binary and ternary PlTasA complexes reveals conformational rearrangements broadly analogous to those in SpTasH, including repositioning of the coiled-coil and Nop domains⁵. In addition, the coiled-coil helices undergo a lateral displacement upon target binding, disrupting the face-to-face alignment seen in the binary complex. This displacement repositions the Nop domain toward the junction, where the conserved α4 pair separates the two DNA strands. Notably, these rearrangements almost exclusively affect protomer B, while protomer A and the tigRNA stem remain fixed relative to each other (**Supplementary Fig. 3**). This asymmetry led us to hypothesize that the two spacers contribute unequally to target engagement, which we tested by probing each independently. Together, these structural and functional analyses establish PlTasA as a compact, nuclease-deficient TIGR-Tas effector that preserves the conserved target-recognition architecture while incorporating structural adaptations to accommodate recognition of a stem-loop tigRNA.

### Asymmetric target engagement by TIGR-Tas systems

The observation of asymmetric rearrangement between the protomers raised the question of whether the two RNA-DNA heteroduplexes contribute equivalently to DNA engagement by PlTasA. To test this, we titrated PlTasA RNP with a fixed concentration of 6-FAM-labeled target DNA and measured binding by electrophoretic mobility shift assay (EMSA) after two hours. Each DNA substrate contained a three-nucleotide bubble positioned within either the Spacer A-or Spacer B-matching region at proximal, medial, or distal locations relative to each unwinding junction (**Fig. 2a**). At the medial and distal positions, Spacer A and Spacer B substrates behaved similarly, with binding remaining in the stoichiometric regime and unsuitable for quantification of dissociation constant (K_d_) values. Strikingly, at the proximal position, the Spacer A substrate reached full engagement, while the Spacer B substrate plateaued near 30–40% at equivalent concentrations (**Fig. 2b**). These results indicate that the two spacer regions contribute asymmetrically to target engagement near the unwinding junctions. This endpoint measurement, however, does not resolve whether the difference is kinetic or reflects a difference in affinity.

**Figure 2.**
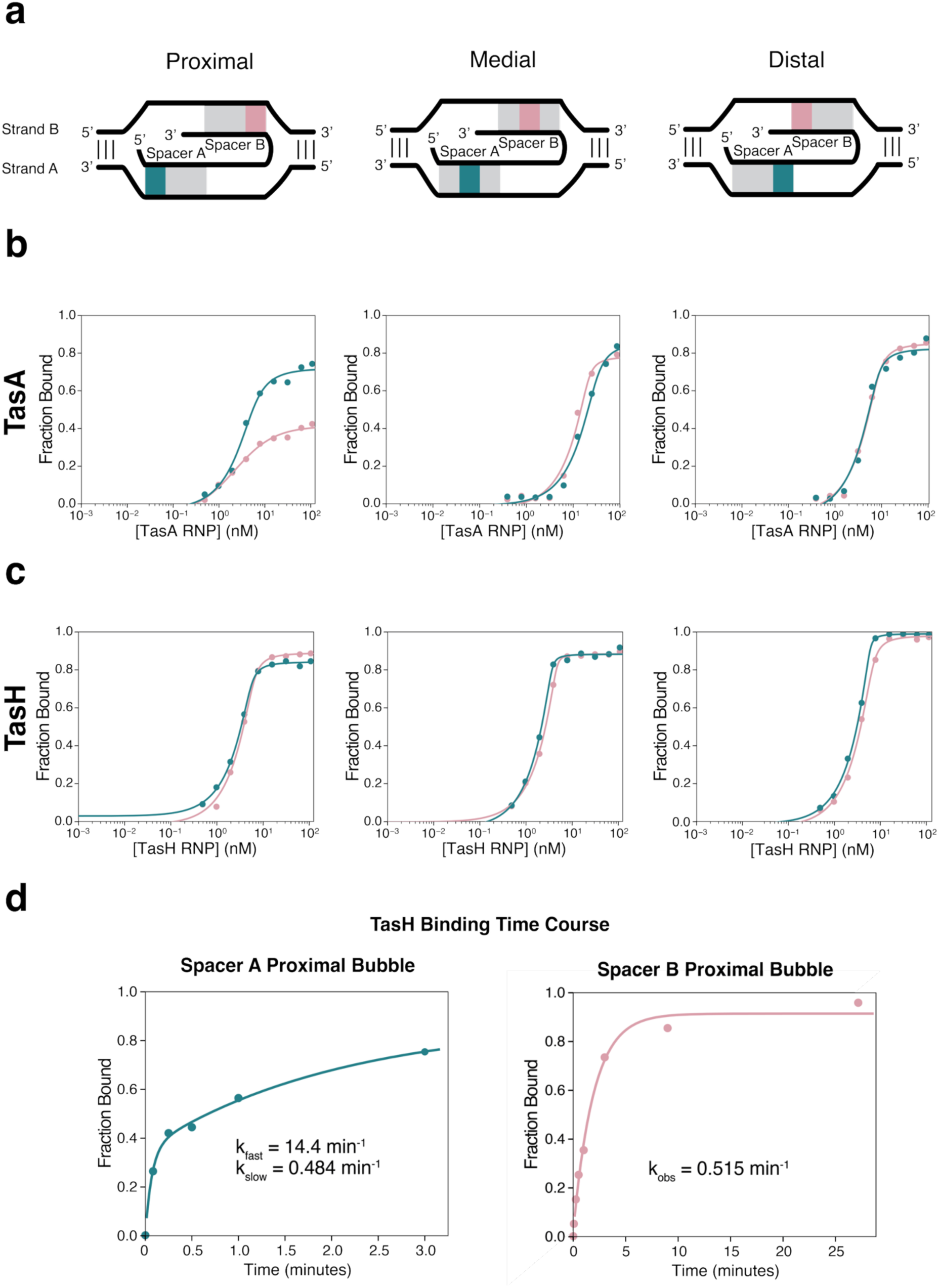
PlTasA and SpTasH both exhibit asymmetrical binding to the target. **a.** Schematic of proximal, medial, and distal bubble placement for each spacer where the teal and salmon colors represent bubble placement at equivalent positions at Spacer A and Spacer B, respectively. **b.** Fraction bound versus PlTasA RNP concentration for proximal (left), medial (center), and distal (right) 6-FAM-labeled DNA bubble substrates, with the bubble positioned within the Spacer A (teal) or Spacer B (salmon) region. Data were fit to a quadratic binding equation. Apparent K_d_ values were not determined, as binding was too tight relative to the substrate concentration to define the titration regime reliably. **c.** Fraction bound versus SpTasH RNP concentration for proximal (left), medial (center), and distal (right) 6-FAM-labeled DNA bubble substrates, with the bubble positioned within the Spacer A (teal) or Spacer B (salmon) region. Data were fit to a quadratic binding equation. Apparent K_d_ values were not determined, as binding was too tight relative to the substrate concentration to define the titration regime reliably. **d.** Time-course binding assays for SpTasH with a proximal DNA bubble positioned within the spacer A (left) or spacer B (right) region. 50 nM SpTasH RNP was used against 5 nM 6-FAM-labeled substrate. Data were fit to a double exponential for spacer A and a single exponential for spacer B, and the derived rates are shown.

To determine whether this asymmetry is conserved across TIGR-Tas effectors, we performed the same assay for SpTasH using the catalytically inactive mutant (H54A), eliminating interference from DNA cleavage. The binding isotherms for each position of both spacers were indistinguishable and remained in the stoichiometric regime (**Fig. 2c**). Because endpoint measurements may conceal kinetic asymmetry, we next monitored binding over time for the proximal-bubble substrates (**Fig. 2d**). Binding to the Spacer A substrate was biphasic, with fast (14.4 min⁻¹) and slow (0.484 min⁻¹) phases, whereas the Spacer B substrate binding was monophasic with a rate constant 0.515 min⁻¹. The similarity between the Spacer A slow phase rate and the single Spacer B rate suggests a shared slow phase, with an additional fast phase unique to Spacer A. Together, these results indicate that the two spacers engage the target asymmetrically in both PlTasA and SpTasH, with Spacer A consistently engaging more rapidly than Spacer B. This suggests that the two RNA-DNA heteroduplexes form in a preferred order.

### SpTasH couples dual-heteroduplex recognition to cleave DNA

To determine how asymmetric DNA engagement affects cleavage by SpTasH, we measured the kinetics of cleavage for both DNA strands. First, SpTasH was assembled with its tigRNA, after which cleavage was initiated by addition of 6-FAM labeled DNA substrate, and quenched at various time points. We observed that SpTasH cleaves both DNA strands at a rate of 0.01 min^−1^ (**Fig. 3a**), approximately 6,000-fold slower than the observed rate for SpCas9 ^9^. To distinguish the cleavage step from preceding target-binding and RNA-DNA heteroduplex formation steps, the DNA substrate was pre-incubated with RNP in the absence of Mg^2+^, followed by the addition of MgCl_2_ to initiate cleavage. This approach bypasses DNA binding and RNA-DNA heteroduplex formation, allowing for measurement of the intrinsic rate constants for DNA cleavage. Under these conditions, we observed a rate of 0.05 or 0.06 min^−1^ for Strand A and Strand B, respectively (**Fig. 3b**). These modest increases indicate that both cleavage and RNA-DNA heteroduplex formation are relatively slow processes and contribute substantially to the overall reaction rate.

**Figure 3.**
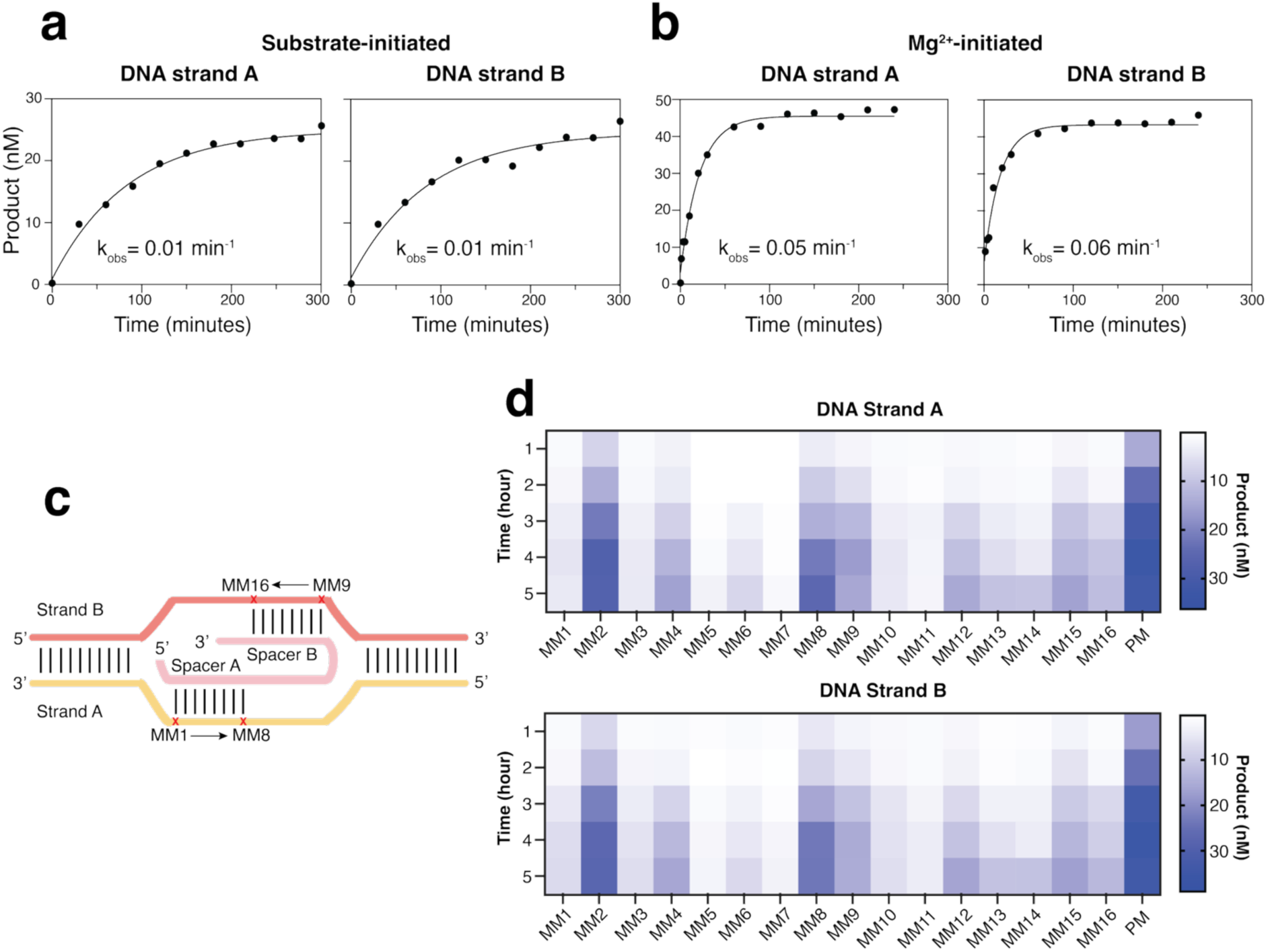
SpTasH exhibits high mismatch sensitivity with DNA cleavage dependent on coordinated recognition of both strands. **a.** Kinetic analyses of SpTasH measured by substrate-initiated cleavage. **b.** Kinetic analyses of SpTasH measured by Mg^2+^-initiated cleavage. **c.** Schematic of the single nucleotide mismatch design. **d.** Heat map shows that DNA cleavage kinetics of Strand A and B with mismatch and perfectly matched DNA (PM).

We next examined the plasmid targeting activity using a GFP reporter assay in *E.coli* as previously described^10^. Upon induction, SpTasH forms a complex with a tigRNA that targets the GFP-encoding plasmid resulting in fluorescence depletion (**Supplementary Fig. 4**). We hypothesized that DNA binding by SpTasH alone might be sufficient to suppress transcription. Consistent with this idea, targeting a nuclease-dead SpTasH mutant (H54A) to either the promoter region or the open reading frame (ORF) of GFP resulted in a reduction in GFP fluorescence (**Supplementary Fig. 4**). Together, these finding establish that SpTasH is an active RNA-guided nuclease both in *in vitro* and in cells. Moreover, DNA binding by a catalytically inactive variant is sufficient to repress transcription, highlighting its potential as a programmable transcriptional repressor.

A previous study demonstrated that a high-fidelity variant of SpCas9 enhances target specificity by reducing the observed cleavage rate, potentially allowing dissociation from the off-target DNA^11^. Given that SpTasH exhibits a markedly slower cleavage rate, we speculated that it may exhibit high target specificity. To test this, we designed a series of single-nucleotide mismatch substrates (MM1-MM16) and compared the cleavage kinetics to that of the perfectly matched (PM) DNA (**Fig. 3c-d**). Overall, SpTasH displayed pronounced sensitivity to target mismatches, with most substitutions strongly suppressing cleavage and appreciable tolerance observed only at positions MM2 and MM8. (**Fig. 3d**). To further assess the mismatch tolerance, we introduced adjacent double mismatches, which abolished cleavage, further supporting the high target specificity of SpTasH (**Supplementary Fig. 5**).

Notably, single mismatches affected cleavage of Strand A and Strand B similarly, despite perturbing only one RNA-DNA heteroduplex, while leaving the opposite strand fully complementary to the tigRNA. Because each DNA strand is cleaved by separate protomers, this concordant response indicates that cleavage at each strand is not governed solely by the local heteroduplex but instead requires coordinated recognition of both spacer-target duplexes. Together with the kinetic observation that the two DNA strands are cleaved at similar rates, these findings support a model in which SpTasH couples dual-heteroduplex surveillance to synchronized DNA cleavage.

### Cryo-electron microscopy structures reveal DNA targeting and activation mechanisms of SpTasH

To determine the structural basis of target engagement, we performed kinetic-informed cryo-EM studies of the SpTasH ternary complex. Guided by the observed kinetics, we incubated target DNA with pre-assembled SpTasH RNP for defined lengths of time before plunge freezing for cryo-EM, with the goal of capturing transient structural intermediates along the target recognition pathway^12^.We first determined a 3.24 Å resolution cryo-EM structure from a dataset in which DNA was incubated with RNP for 30 minutes (**Fig. 4a,b**). This structure shows that the 8 nt Spacer A of tigRNA base-pairs with the DNA strand A, where the complementary region in DNA strand B is displaced. By contrast, Spacer B and the remaining regions of both DNA strands, were not modeled because of weak and diffuse map density, indicating high flexibility and suggesting the heteroduplex between Spacer B and Strand B may not yet be formed. This structural intermediate is consistent with target engagement proceeding in a preferred order, beginning with Spacer A.

**Figure 4.**
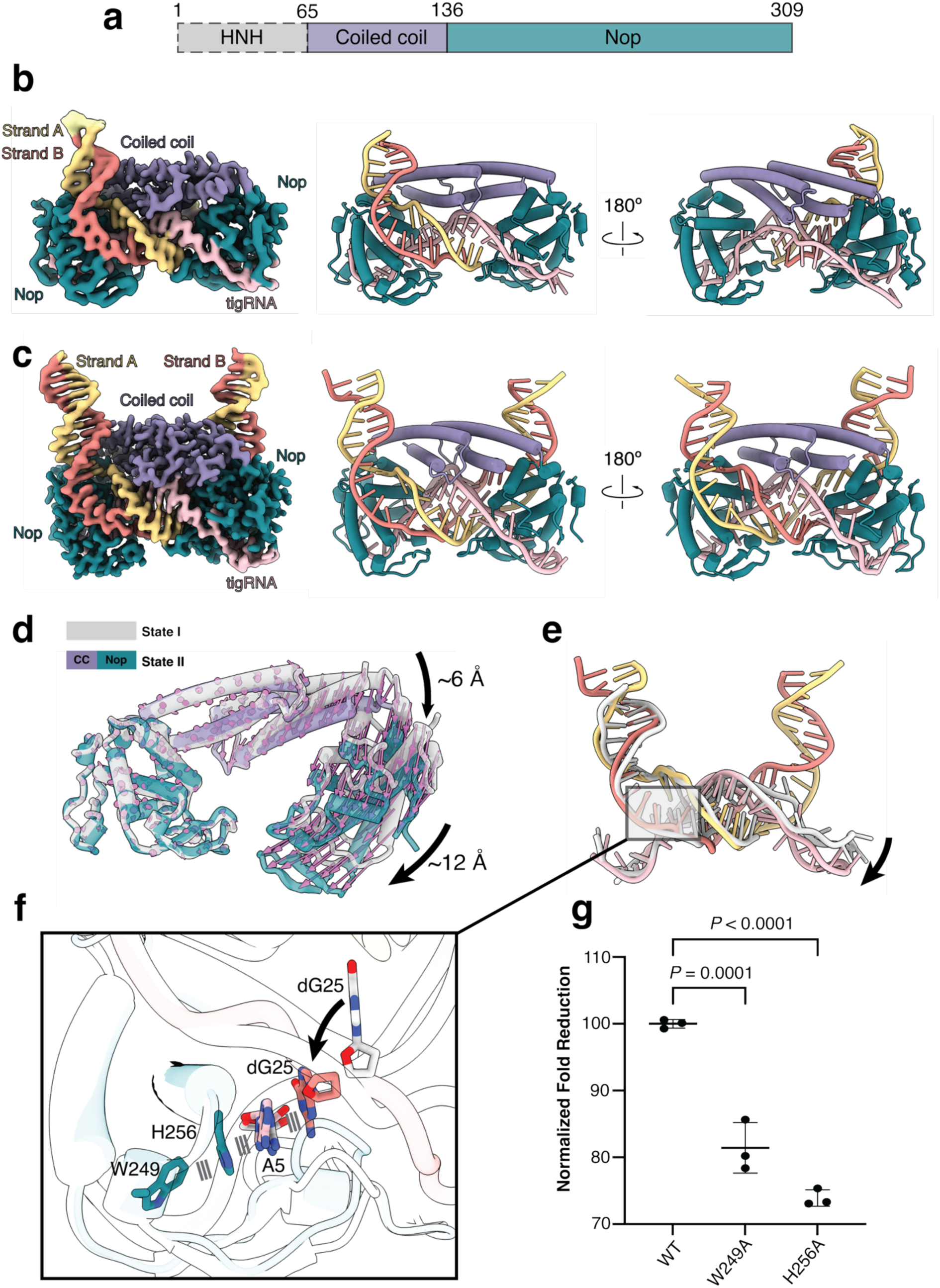
Cryo-EM structures of SpTasH State I and State II reveal distinct stages of RNA-DNA heteroduplex formation. **a.** Domain organization of SpTasH. The HNH domain is colored gray, while the coiled-coil and Nop domains are colored purple, and teal, respectively. **b.** EMReady2-processed cryo-EM map of the SpTasH ternary complex at State I (left), and structural model (middle and right). Map and model are colored based on proximity to the modeled domains shown in **a**. The tigRNA is colored pink, while DNA strands A and B are colored gold and orange, respectively. **c.** EMReady2-processed cryo-EM map of the SpTasH ternary complex at State II (left), and structural model (middle and right). Map and model are colored based on proximity to the modeled domains shown in a. The tigRNA is colored pink, while DNA strand A and B are colored gold and orange, respectively. **d.** Structural alignment of State I and State II that shows conformational changes in protomer B. Arrows between models were generated using PDBarrows v.1.0 as described in Chaaban et al^29^, showing domain movements from State I to State II. **e.** Structural alignment of the tigRNA in State I and State II. **f.** Zoom in figure shows Spacer B base pairing induced the proper interaction of the dG25 of Strand B with tigRNA and protomer A in a conserved base-stacking interaction. **g.** GFP depletion activity of SpTasH wild-type and mutants. Data represent mean ± SD (n = 3 biological replicates). Significance was determined by one-way ANOVA.

We next determined a second structure, collected after 2 hours of DNA incubation with RNP, of the fully engaged ternary complex at a resolution of 2.34 Å. This structure reveals both Spacer A and Spacer B are base-paired with the complementary DNA, indicating that prolonged incubation allows full formation of the RNA-DNA heteroduplex (**Fig. 4c**). SpTasH forms a C2-symmetric homodimer, with the coiled-coil domain serving as the dimerization interface (**Supplementary Fig. 6**). Each coiled-coil domain contains two long α-helices that pack against the corresponding helices of the other protomer, forming a two-layer architecture. The top layer dimer interface is stabilized by a cation-π interaction between F109 and R120, as well as hydrogen-bonding interactions involving R120, S105 and R103 (**Supplementary Fig. 6**). Alanine substitutions of these residues resulted in a defect in plasmid targeting relative to the WT in the GFP depletion assay (**Supplementary Fig. 6**), underscoring the essential role of dimer stability in function. Unlike PAM recognition in CRISPR-Cas systems which involves sequence-specific interactions with the DNA bases, for SpTasH, we observed sequence-independent interactions between basic residues (R213, R218, K225) and the phosphate backbone of Strand B (**Supplementary Fig. 6**). This observation is consistent with the fact that SpTasH does not have PAM requirement. Alanine substitution mutants displayed reduced plasmid targeting ability compared to the WT (**Supplementary Fig. 6**), suggesting disruption of this protein-DNA interactions impairs SpTasH function.

Structural alignment of State I and State II reveals that formation of the Spacer B - Strand B duplex drives conformational changes both in SpTasH and the tigRNA (**Fig. 4d**). While protomer A remains largely unchanged, protomer B undergoes substantial rearrangements, including a ∼6 Å shift of the coiled-coil domain and a ∼12 Å downward displacement of the Nop domain (**Fig. 4d**), These movements accommodate formation of the Spacer B-Strand B heteroduplex. Accordingly, the tigRNA also shifts downward to stabilize the complex (**Fig. 4e**). Additionally, formation of Spacer B-Strand B heteroduplex affects the arrangement of the displaced Strand B adjacent to the Spacer A-Strand A heteroduplex. Specifically, one nucleotide, dG25, undergoes a pronounced repositioning and interacts with the A5 of the tigRNA, forming an extended base-stacking interaction together with the residues H256 and W249 of the protomer A (**Fig. 4f**). Disruption of this interaction by mutating either H256 or W249 decreased the plasmid targeting ability of SpTasH (**Fig. 4g**), suggesting Spacer B pairing also influences the stability and interaction between DNA, tigRNA and effector, thereby contributing to coordinated target recognition and efficient DNA cleavage.

### TIGR–TasA–VPR activates GFP expression in HEK293T cells

Based on the structural and biochemical characterization of TasA-mediated DNA recognition, we sought to establish TasA as a programmable transcriptional regulator in mammalian cells by fusing FpTasA and PlTasA to the VPR activation domain^13^. To evaluate TasA-VPR-mediated transcriptional activation, an SV40 promoter-driven GFP reporter was used together with two tigRNAs (SV40-T1 and SV40-T2) targeting distinct sites within the SV40 promoter, while a non-targeting tigRNA (NT) served as the control (**Fig. 5a**). The fluorescence microscopy and flow cytometry results showed that FpTasA-VPR increased GFP expression when guided by SV40-T1 tigRNA, whereas no significant activation was observed with SV40-T2. In contrast, significant GFP activation by PlTasA-VPR was observed with both tigRNAs (**Fig. 5b; Supplementary Fig. 11-12**). To further validates the transcriptional activation, the remaining cells from flow cytometry were collected for RNA extraction and RT-qPCR analysis. RT-qPCR showed significant GFP mRNA upregulation in the same groups identified as significantly activated by flow cytometry (**Fig. 5c).** These results demonstrate that both FpTasA-VPR and PlTasA-VPR can mediate programmable transcriptional activation in mammalian cells.

**Figure 5.**
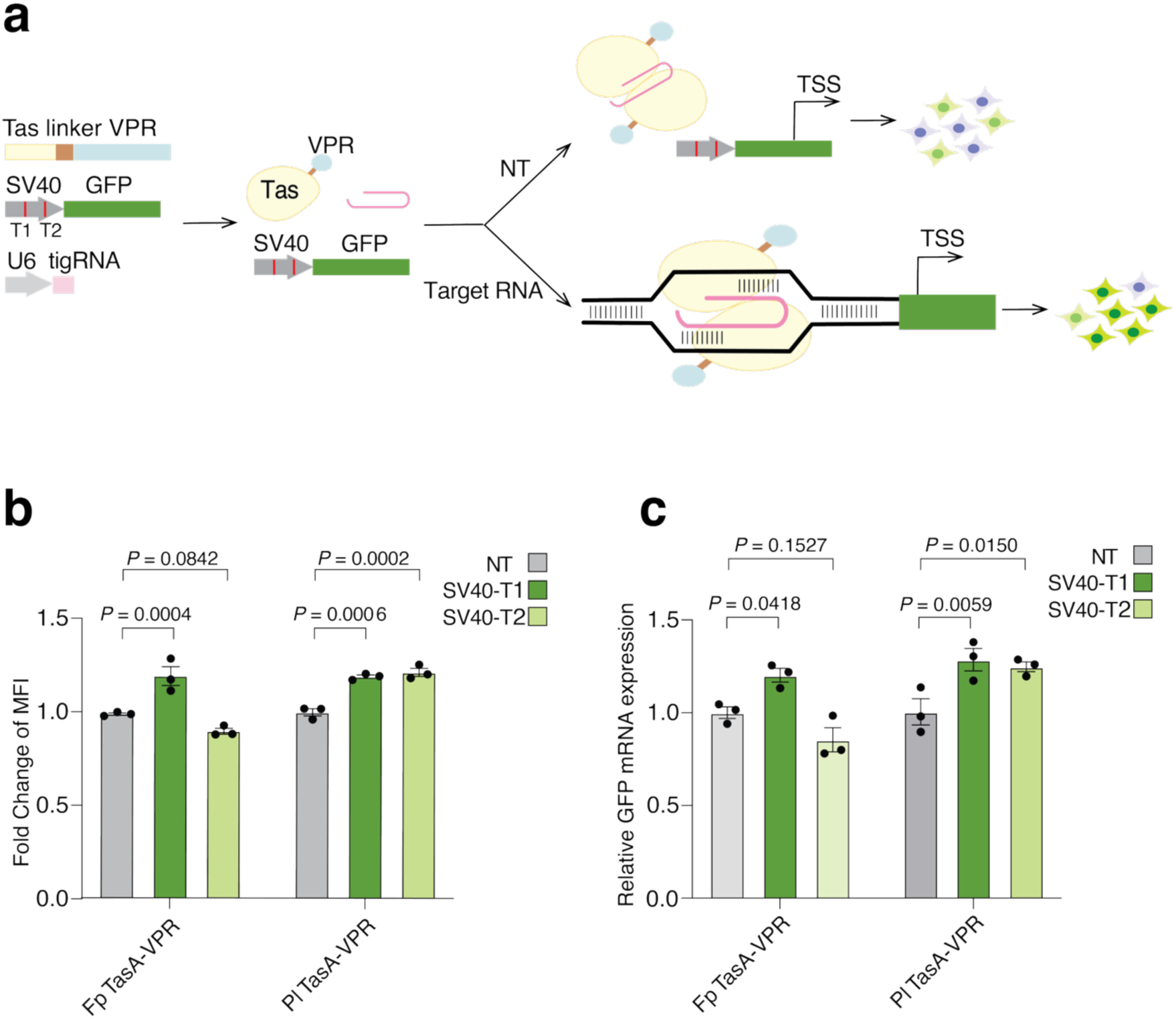
TIGR-TasA-VPR-mediated transcriptional activation of GFP in mammalian cells. **a.** Schematic of the TasA-VPR transcriptional activation system in HEK293T cells. Cells were co-transfected with an SV40 promoter-driven GFP reporter, TasA-VPR, and a U6 promoter-driven tigRNA (pink) containing a non-targeting (NT), SV40-T1, or SV40-T2 guide. FpTasA or PlTasA (yellow) was fused to the VPR activation domain (cyan) to activate GFP transcription. tigSV40-T1 and tigSV40-T2 target the SV40 promoter, with their target sites indicated in red. **b.** GFP protein activation quantified by flow cytometry as mean fluorescence intensity (MFI). MFI values are shown as fold change relative to the corresponding NT control. **c.** GFP transcriptional activation quantified by RT-qPCR and normalized to GAPDH. In both experiments, FpTasA-VPR or PlTasA-VPR was co-transfected with NT, SV40-T1, or SV40-T2 tigRNA plamids. Error bars represent mean ± SEM from 3 independent biological replicates. Statistical significance was determined by two-way ANOVA followed by Šídák’s multiple-comparisons test.

## Discussion

TIGR-Tas systems expand the repertoire of RNA-guided effectors through their tandem-spacer guide architecture, compact size, and apparent lack of a PAM requirement. Here, we show that these features are accompanied by an ordered mechanism of target recognition that is conserved across catalytically distinct TIGR-Tas effectors. In this study, we characterized *Peromyscus leucopus* TasA (PlTasA), a nuclease-deficient effector, and *Salicola phage* TasH (SpTasH), an RNA-guided DNA nuclease. Structural analysis showed that PlTasA preserves the core architecture of previously characterized TIGR-Tas effectors despite lacking a nuclease domain and associating with a stem-loop rather than a dual-repeat tigRNA (**Fig. 1b-d**). Residues corresponding to the aromatic-basic pair implicated in DNA unwinding are preserved, indicating that target recognition and duplex separation follow a common mechanism across TIGR-Tas systems. Additionally, PlTasA contains contacts that accommodate a stem-loop tigRNA while maintaining the spatial arrangement of the two spacer-target heteroduplexes (**Fig. 1e)**.

Comparison of the PlTasA binary and ternary complexes reveals that Protomer A largely retains its position relative to the tigRNA, whereas protomer B undergoes a substantial rearrangement upon target binding (**Supplementary Fig. 3**). This organization is consistent with a directional model in which protomer A is preorganized for initial DNA recognition, while protomer B subsequently rearranges to accommodate formation of the Spacer B heteroduplex. The two SpTasH ternary structures support a similar stepwise mechanism. The earlier state contains only the Spacer A-Strand A heteroduplex, whereas prolonged incubation with the DNA allowed formation of both heteroduplexes and induced substantial rearrangements in protomer B (**Fig. 3b-d**). Spacer B pairing also repositions dG25 of Strand B into a stacking network with tigRNA nucleotide A5 and residues H256 and W249 of protomer A (**Fig. 4f**). Together with the previously observed requirement for A5 in Strand A cleavage^5^, these findings indicate that formation of the second heteroduplex stabilizes interactions across the complex and promotes coordinated target recognition.

Our binding measurements further demonstrate that the two tigRNA spacers engage the target asymmetrically. A proximal DNA bubble affects target association differently depending on whether it was introduced within the Spacer A-or Spacer B-matching region (**Fig. 2a-d**). Because this was observed in both PlTasA and SpTasH, which differ in nuclease content and guide architecture, asymmetric target engagement is likely a general feature of TIGR-Tas systems. Interestingly, a recent structure of SpTasH bound to a target containing a mismatch at position 5 of the Spacer A (MM5) showed that Spacer A-target heteroduplex was unresolved, whereas the Spacer B-target heteroduplex remained fully formed^14^. Although this observation is inconsistent with a strictly ordered pathway in which Spacer A must engage before stable formation of the Spacer B heteroduplex, it is compatible with a preferred order of engagement. We propose that Spacer A normally engages before stable formation of the Spacer B heteroduplex, but that a strongly destabilizing mismatch can disrupt this preference and allow Spacer B pairing to proceed independently, thereby trapping the complex in an alternative partially engaged state.

Unlike canonical CRISPR-Cas nucleases, which typically show sequential DNA strand cleavage^9^, our results indicate that SpTasH cleaves both DNA strands with indistinguishable kinetics, albeit with a relatively low rate (**Fig. 3a-b**). This observation is consistent with a mechanism that promotes fidelity by allowing dissociation of off-target DNA. Indeed, most single mismatches strongly reduced cleavage by SpTasH, and adjacent double mismatches abolished detectable activity (**Fig. 3d; Supplementary Fig. 5**). Notably, individual mismatches affected cleavage of both DNA strands similarly indicating that the two catalytic centers are functionally coupled and that productive cleavage requires coordinated surveillance of both spacer-target duplexes.

Intriguingly, our mismatch-profiling results differ from those reported in a recent study showing a mismatch at the first position of Spacer A (MM1) was largely tolerated, whereas a mismatch at position 5 (MM5) converted SpTasH into a strand-specific nickase^14^. In contrast, most single mismatches impaired cleavage of both DNA strands to a similar extent, supporting coordinated surveillance of the two heteroduplexes. The apparent discrepancy may arise from differences in guide-target sequences, metal-ion concentrations, reaction conditions, or the time points used to evaluate cleavage, like what has been shown previously that spacer sequence and metal ion concentration can affect the mismatch tolerance of CRISPR-Cas enzymes^15,16^. Thus, certain mismatches may permit partial assembly and asymmetric HNH recruitment, whereas most mismatches prevent the coordinated conformational maturation required for efficient cleavage of either strand. Further kinetic and structural analyses using the same guide sequences and reaction conditions will be required to determine how sequence context governs the balance between coordinated double-strand cleavage and mismatch-induced nicking.

Collectively, our results support a conserved and ordered mechanism of TIGR-Tas target recognition. Spacer A preferentially initiates target engagement; Spacer B pairing promotes structural maturation and stabilizes interactions across the complex. Completion of both heteroduplexes enables coordinated effector activity. In SpTasH, this pathway produces slow DNA cleavage with indistinguishable strand-specific kinetics and pronounced mismatch sensitivity, whereas in PlTasA the conserved recognition scaffold supports stable DNA binding. These properties establish TIGR-Tas systems as attractive platforms for developing compact, PAM-independent tools for genome editing and transcriptional regulation.

## Materials and methods

### Plasmid construction

The codon-optimized coding sequences of SpTasH and PlTasA were synthesized by Integrated DNA Technologies (IDT) and cloned into a pET-29b expression vector with a C-terminal 6xHis tag by PCR using Q5 High-Fidelity DNA Polymerase and Gibson Assembly Master Mix (NEB). Site-directed mutagenesis was carried out using Q5 High-Fidelity DNA Polymerase and KLD Enzyme Mix (NEB). All constructs were verified by Oxford Nanopore Technology long-read sequencing (Plasmidsaurus). For mammalian cell transfection, NLS-containing, mammalian codon-optimized FpTasA and PlTasA sequences were synthesized by Twist Bioscience. The synthesized TasA sequences, together with the linker and VPR sequences derived from NLS-dCasMINI-V4-VPR-c-Myc_NLS-mCherry-WPRE (Addgene #176269), were cloned by HiFi DNA Assembly (NEB, E2621S) into the backbone of the previously reported AsCas12a-P2A-GFP plasmid, replacing the original coding sequence^17^. The FpTasA and PlTasA sequences were also cloned downstream of the U6 promoter in U6_crRNA_mCherry (Addgene #197609) by HiFi DNA Assembly (NEB, E2621S), replacing the Cas7-11 crRNA scaffold. The reporter plasmid was obtained from Addgene (plasmid #46919), a gift from Arbor Biotechnologies.

### Nucleic acid preparation

tigRNA and target DNA oligos were synthesized by Integrated DNA Technologies (IDT). The sequences, including the positions of mismatches in DNA, are listed in **Supplementary Data 1**.

### Protein production and purification

SpTasH and PlTasA were expressed and purified as previously described with minor modifications^4^. Briefly, protein expression was induced overnight in *Escherichia coli* Nico21(DE3) (NEB) for SpTasH and BL21(DE3) (NEB) for PlTasA with 0.5 mM IPTG at 16 °C. Cells were harvested by centrifugation and lysed by sonication in Lysis buffer (20 mM HEPES pH 7.5, 500 mM NaCl, 10 mM imidazole). The lysate was clarified by centrifugation at 38,000 x g for 45 minutes at 4 °C. The supernatant was loaded onto a HisTrap HP column (Cytiva) for affinity purification. Protein was eluted with an imidazole gradient buffer. The collected eluate was concentrated and loaded onto a Superdex 200 Increase 10/300 column (Cytiva) pre-equilibrated with SEC buffer (20 mM HEPES pH 7.5, 200 mM NaCl, 0.5 mM TCEP and 10 mM MgCl_2_). Peak fractions were collected and examined by SDS-PAGE. Protein was flash-frozen in liquid nitrogen and stored at -80 °C.

### Cryo-EM sample preparation, data collection and processing

Prior to complex assembly, tigRNA and dsDNA (equimolar targeting and non-targeting strand) were prepared in annealing buffer (10 mM Tris-HCl, pH 7.5, 50 mM NaCl, 1 mM DTT), followed by heating at 95 °C for 5 min and cooling at room temperature for 15 min. SpTasH-tigRNA and PlTasA-tigRNA binary complexes were assembled by mixing purified Tas protein with its cognate tigRNA at a molar ratio of 1:1.5 followed by incubation at 37 °C for 30 min. For the ternary complexes, equimolar dsDNA duplex was then incubated with the binary complex at 37 °C for 30 min (SpTasH State I), or 2 hour (SpTasH State II and PlTasA).

For grid preparation, 2.5 μL 7 μM sample was applied onto the Quantifoil R1.2/1.3 400 mesh copper grid that had been plasma-cleaned for 30 s by a Solarus 950 plasma cleaner (Gatan). Grids were blotted by a Vitrobot Mark IV (Thermo Fisher) for 8 s with blot force of 0 at 4 °C and 100% humidity, and plunge-frozen in liquid ethane. Grids were stored in liquid nitrogen before screening.

For the PlTasA binary complex, dataset was collected with SerialEM^18^ on a FEI Glacios cryo-TEM equipped with a Falcon 4 detector with a pixel size of 0.933 Å. The defocus range was set to –1.5 to –2.5 μm. 3,411 accepted micrographs were processed by motion correction, contrast transfer function (CTF) estimation in cryoSPARC live^19^ . Picking was done by blob picker with a particle diameter range of 60 - 160 A °. 3,713,229 particles were extracted at a box size of 320 pixels with a Fourier crop to 128 pixels and then classified into 50 2D classes. 1,459,107 particles were selected from 2D classification for further ab-initio reconstruction (3 classes). The resulting particles and volume classes were subjected to heterogeneous refinement. Iterative ab-initio followed by heterogenous refinement was performed three total times. The best-resolved class was selected for particle re-extraction at box size of 320 pixels followed by non-uniform refinement which yielded a volume at 4.09 Å resolution. The particles were further processed by reference-based motion correction and a final round of non-uniform refinement which yielded a volume at 3.90 Å (**Supplementary Fig. 7,8**).

For the PlTasA ternary complex, datasets were collected with SerialEM^18^ on a FEI Krios cryo-TEM equipped with a Gatan K3 direct electron detector with a pixel size of 0.8332 Å. The defocus range was set to –1.5 to –2.5 μm. 4,512 accepted micrographs were processed by motion correction, contrast transfer function (CTF) estimation in cryoSPARC live^19^. Picking was done by blob picker with a particle diameter range of 60 - 160 A °. 4,843,453 particles were extracted at a box size of 320 pixels with a Fourier crop to 128 pixels and then classified into 50 2D classes. 836,787 particles were selected from 2D classification for further ab-initio reconstruction (3 classes). The resulting particles and volume classes were subjected to heterogeneous refinement. The best-resolved class was selected for particle re-extraction at box size of 320 pixels followed by non-uniform refinement which yielded a volume at 3.18 Å resolution. The particles were further processed by Local and Global CTF refinement, reference-based motion correction, and a final round of non-uniform refinement which yielded a volume at 3.06 Å (**Supplementary Fig. 7,8**). The final map was processed by Ardecon 1.0^20^ to address the preferred orientation issue and then EMReady2^21^.

For SpTasH State I, dataset was collected with SerialEM^18^ on a FEI Glacios cryo-TEM equipped with a Falcon 4 detector with a pixel size of 0.933 Å. The defocus range was set to – 1.5 to –2.5 μm. 2, 498 accepted micrographs were processed by motion correction, contrast transfer function (CTF) estimation in cryoSPARC live^19^. Picking was done by blob picker with a particle diameter range of 60 - 120 A °. 2,527,331 particles were extracted at a box size of 320 pixels with a Fourier crop to 128 pixels and then classified into 50 2D classes. 2D classification was performed using a maximum resolution of 8 A °, 500 batch size and 40 online-EM iterations, all other settings were kept at default. 629,897 particles were selected from 2D classification for further ab-initio reconstruction (3 classes). The resulting particles and volume classes were subjected to heterogeneous refinement. The best-resolved class was selected for particle re-extraction at box size of 256 pixels. followed by non-uniform refinement which yield a volume at 3.55 Å resolution. The particles were further processed by reference-based motion correction and the final round of non-uniform refinement which yield a volume at 3.24 Å that used for modeling (**Supplementary Fig. 9,10**). The final map was processed by EMReady2^21^.

For SpTasH State II, dataset was collected with SerialEM^18^ on a FEI Krios cryo-TEM equipped with a Gatan K3 direct electron detector with a pixel size of 0.8332 Å. The defocus range was set to –1.5 to –2.5 μm. 3,880 accepted micrographs were processed by MotionCor2^22^, then imported to cryoSPARC (v4.7.1)^19^ for Patch CTF estimation, as well as blob picking with a particle diameter range of 60 - 120 A °. 4,989,309 particles were extracted at a box size of 320 pixels with a Fourier crop to 128 pixels and then classified into 50 2D classes. 3,198,004 particles were selected from 2D classification for further ab-initio reconstruction (3 classes). The resulting particles and volume classes were subjected to heterogeneous refinement. The best-resolved class was selected for particle re-extraction at box size of 300 pixels. followed by non-uniform refinement which yield a volume at 2.34 Å resolution that was used for modeling (**Supplementary Fig. 9,10**). The final map was processed by EMReady2^21^.

### Model building and refinement

The ternary complex structure predicted by AlphaFold3^23^ was fitted into the map as a rigid body in ChimeraX (v1.9)^24^. The model was manually adjusted in Coot (v0.9.8.95 EL)^25^ and automatically refined by real_space_refine in Phenix (v2.0-5936)^26^. All structural figures were generated using ChimeraX (v1.9).

### Electrophoretic mobility shift assays

Binding of PlTasA and SpTasH RNP complexes to DNA was assessed by electrophoretic mobility shift assay (EMSA). tigRNA and 6-FAM-labeled DNA duplexes were prepared by denaturing at 95 °C for 5 min and cooling to room temperature over 15 min in annealing buffer (20 mM HEPES pH 7.5, 120 mM NaCl, 0.5 mM TCEP). RNPs were assembled by incubating purified PlTasA with cognate tigRNA at a 1:1.2 molar ratio in binding buffer (20 mM HEPES pH 7.5, 150 mM NaCl, 10 mM MgCl₂, 0.5 mM TCEP, 0.05% Tween-20) at 37 °C for 40 min; PlTasA concentrations refer to the dimer unless otherwise stated. Assembled RNP was combined with 6-FAM-labeled dsDNA (5 nM final) in binding buffer (10 μL final volume) and incubated at 37 °C, either for 2 h (endpoint) or for the indicated times (time course). Reactions were quenched with an equal volume of ice-cold 2× native Tris-glycine loading dye (Invitrogen, LC2673) and held on ice. For time-course reactions, the dye was supplemented with a 50-fold molar excess of the corresponding unlabeled DNA substrate to prevent rebinding. Samples were resolved on 6% Tris-glycine polyacrylamide gels (Invitrogen, XP00065BOX) pre-run for 90 min at 100 V, 4 °C, and run at 100 V for 45 min at 4 °C in Tris-glycine buffer. 6-FAM fluorescence was imaged on an Amersham Typhoon RGB scanner (Cytiva) and quantified by densitometry in Fiji ^27^. Fraction bound was calculated as bound signal divided by total signal per lane, and apparent binding was compared qualitatively across conditions without determining dissociation constants. The time-resolved assays were fit to single-or double-exponential equations in KinTek Explorer ^27^ (v2026.1.10).

### GFP depletion assay

GFP depletion assays were performed as previously described^10^. SpTasH WT or mutants coding sequence and DNA template of tigRNA were cloned into *araBAD*-driven plasmids with different antibiotic resistance. The GFP-encoding plasmid was used as the target. 5 μL of *E. coli* DH5α overnight culture bearing these three plasmids were sub-cultured to 2.5 mL fresh LB medium. The culture was then split into two equal volumes: (1) as an uninduced control and (2) induced with 20 mM L-arabinose. All cultures were distributed into a 96-well plate (Invitrogen), then incubated at 37 °C at 300 rpm. GFP fluorescence was monitored overnight by a CLARIOstar Plus plate reader (BMG Labtech). Each measurement was done at 3 independent biological replicates. Fold reduction was calculated by comparing fluorescence between control and induced samples.

### DNA cleavage kinetics

For substrate-initiated cleavage, reactions were initiated by adding 50 nM 6-FAM labeled DNA substrate into pre-assembled 500 nM SpTasH-tigRNA complex in reaction buffer (20 mM Tris-HCl, pH 7.5, 100 mM KCl, 5% glycerol, 10 mM MgCl_2_, 1mM DTT). Reactions were performed at 37 °C then quenched at various times by mixing with equal volume of 0.5 M EDTA. For Mg^2+^-initiated cleavage, 50 nM 6-FAM labeled DNA was incubate with 500 nM SpTasH-tigRNA complex in reaction buffer without MgCl_2_ (20 mM Tris-HCl, pH 7.5, 100 mM KCl, 5% glycerol, 1mM DTT) for 2 hours. Reactions were initiated by adding MgCl_2_ to 10 mM then quenched at various times by mixing with equal volume of 0.5 M EDTA. Reaction products were analyzed by 15% TBE-Urea gel (BioRad). Gels were run in 1X TBE buffer at 180 V for 1 hour and visualized on a ChemiDoc gel imager (BioRad). Fluorescence intensities were quantified using Fiji^27^. For mismatch DNA cleavage kinetics, two DNA strand were labeled with 6-FAM and Cy5, respectively, and DNA cleavage was performed as described above. Data were fit using either a single or double-exponential equations in GraphPad Prism.

### Mammalian cell culture

HEK293T (ATCC #CRL-3216) cells were cultured in DMEM high glucose GlutaMAX^TM^ supplement pyruvate (Gibco #10569010), 10% Fetal Bovine Serum (Gibco #A3160902) and 1X Penicillin-Streptomycin. Cells were incubated at 37°C and 5% CO_2_.

### Mammalian cell transfection

For the SV40-GFP reporter assay, HEK293T cells were seeded at 10^5^ cells per well in 200 µL of complete DMEM in 48-well plates. After 24 h, cells were co-transfected by TransIT-X2® transfection reagent (Mirus Bio, Cat. #000001346) with 400 ng of the SV40-GFP reporter plasmid, 450 ng of FpTasA-VPR or PlTasA-VPR plasmid, and 50 ng of the indicated tigRNA plasmid (NT, SV40-T1, or SV40-T2). 48 h after transfection, cells were imaged by fluorescence microscopy using an EVOS M5000 Imaging System (Thermo Fisher Scientific, AMF5000). Cells were then harvested for flow cytometry analysis and RNA extraction.

### Flow cytometry for quantification of GFP expression

After 48 h transfection, cells were trypsinized with 1× Trypsin-EDTA (Gibco, Cat. #25200-056), resuspended in complete DMEM, and filtered through a 35-µm cell strainer (Falcon, Cat. #352235). Samples were analyzed on a CytoFLEX LX flow cytometer (Beckman Coulter) using the blue laser with a 510/20-nm filter to detect GFP fluorescence. A total of 10,000 events were recorded for each sample. FCS files were analyzed using FlowJo™ v10.10, and the mean fluorescence intensity (MFI) of GFP-positive cells was quantified.

### RNA extraction and RT-qPCR for GFP

Total RNA was extracted from cells using the Maxwell® RSC simplyRNA Cells Kit (Promega, AS1390) on a Maxwell® RSC Instrument (Promega, AS4500), according to the manufacturer’s instructions. RT-qPCR was performed by the Luna® Universal Probe One-Step RT-qPCR Kit (NEB, E3006S) on an Applied Biosystems QuantStudio™ 5 Real-Time PCR System. Reactions were multiplexed using a FAM-labeled probe for GFP and a Cy5-labeled probe for GAPDH, with GAPDH serving as the reference gene. The GAPDH primers and probe were adopted from Asahi-Ozaki et al^28^ and the sequences of the GFP and GAPDH primers and probes used in this study are provided in **Supplementary Data 1**. Relative GFP expression was calculated using the ΔΔCt method.

## Data availability

Structure of the PlTasA-tigRNA binary complex have been deposited in the EMDB with accession codes: EMD-X. Structure of the PlTasA-tigRNA-DNA ternary complex have been deposited in the EMDB with accession codes: EMD-X, Associated atomic coordinates were deposited to PDB with accession code: X. Structures of the SpTasH-tigRNA-dsDNA State I and State II have been deposited in the EMDB with accession codes: EMD-X, and EMD-X, respectively. Associated atomic coordinates were deposited to PDB with accession codes: X , respectively.

## Acknowledgements

We thank Taylor lab members, particularly Drs. Matthew M. Hooper and Rodrigo Fregoso Ocampo, for valuable discussion. We thank Drs. Axel Brilot and Evan Schwartz at the Sauer Lab at UT Austin for cryo-EM assistance. This work was supported by National Institutes of Health grant R35GM138348 (to D.W.T.) and NIH-NIGMS Maximizing Investigator’s Research Award (MIRA) R35GM147788 (P.K.J.).

## Author contributions

K.G. and N.M.A. conceived the experiments; prepared samples for cryo-EM studies; collected and processed Cryo-EM datasets, modeled the structures. N.M.A. and G.S. performed EMSA binding assays. K.G. performed DNA cleavage kinetics. K.G. and K.M.M.A. performed GFP depletion assays. D.C. and B.H. performed Mammalian cell transfection, flow cytometry and RT-qPCR. K.G., N.M.A., B.H. and D.C. wrote the manuscript. P.K.J. and D.W.T. supervised the project, secured funding, and edited the manuscript. All authors reviewed and approved the manuscript.

