## Supplementary Information for "Asymmetric DNA targeting by RNA-guided TIGR-Tas systems"

**Table 1 Cryo-EM data collection, refinement and validation statistics.**

| | **PlTasA binary** | **PlTasA ternary** | **SpTasH-State I** | **SpTasH-State II** |
|---|---|---|---|---|
| **Data collection and** | | | | |
| Magnification | 150,000 | 105,000 | 150,000 | 105,000 |
| Voltage (kV) | 200 | 300 | 200 | 300 |
| Electron exposure (e–/Å$^2$) | 49 | 70 | 49 | 70 |
| Defocus range (μm) | –1.5 to –2.5 | –1.5 to –2.5 | –1.5 to –2.5 | –1.5 to –2.5 |
| Pixel size (Å) | 0.933 | 0.8332 | 0.933 | 0.8332 |
| Initial particle (no.) | 3,713,229 | 4,843,453 | 2,527,331 | 4,989,309 |
| Final particle (no.) | 93,240 | 440,808 | 293,790 | 2,206,371 |
| Map resolution (Å) | 3.9 | 3.06 | 3.24 | 2.34 |
| FSC threshold | 0.143 | 0.143 | 0.143 | 0.143 |
| **Refinement** | | | | |
| Initial model used | | AlphaFold3 | AlphaFold3 | AlphaFold3 |
| Model resolution (Å) | - | 2.9 | 3.6 | 2.7 |
| FSC threshold | - | 0.5 | 0.5 | 0.5 |
| Map sharpening B factor (Å$^2$) | - | –162.8 | –180 | –117.4 |
| Model composition | - | | | |
| Non-hydrogen atoms | - | 5410 | 5196 | 6027 |
| Protein residues | - | 438 | 476 | 478 |
| Nucleotides | - | 90 | 66 | 106 |
| Ligands | - | N/A | N/A | N/A |
| Mean B factors (Å$^2$) | - | | | |
| Protein | - | 61.13 | 87.65 | 53.21 |
| Nucleotides | - | 69.77 | 101.17 | 55.08 |
| R.m.s deviations | - | | | |
| Bond lengths (Å) | - | 0.007 | 0.006 | 0.007 |
| Bond angles (°) | - | 1.308 | 1.296 | 1.299 |
| **Validation** | - | | | |
| MolProbity score | - | 1.53 | 1.64 | 1.29 |
| Clashscore | - | 8.48 | 7.18 | 5.35 |
| Poor rotamers (%) | - | 0.8 | 1.26 | 1.01 |
| Ramachandran plot | - | | | |
| Favored (%) | - | 97.65 | 97.02 | 98.94 |
| Allowed (%) | - | 2.35 | 2.98 | 1.06 |
| Disallowed (%) | - | 0 | 0 | 0 |
| **PDB** | **-** | | | |
| **EMDB** | **EMD-** | **EMD-** | **EMD-** | **EMD-** |

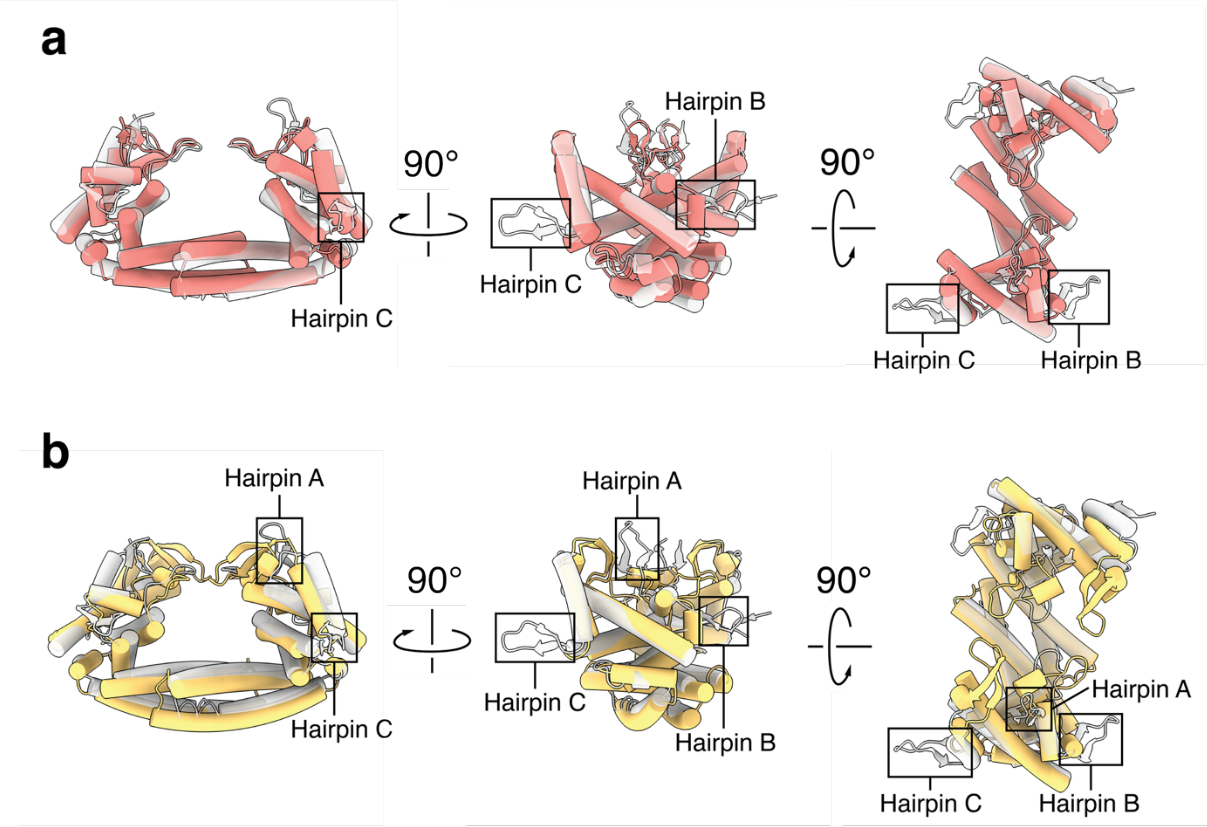


**Supplementary Figure 1. Superposition of PlTasA with TaTasR and SpTasH.**

**a.** PlTasA (grey) superimposed on TaTasR (salmon; PDB: 9MTY), and **b**. PlTasA (grey) superimposed on SpTasH (gold; PDB: 9W04). Superpositions were performed using CC-Nop core (secondary-structure pairing, iteration cutoff 2.0 Å, excluding the HNH and RuvC nuclease insertions absent from PlTasA), giving RMSD values of 1.2 Å over 224 Cα atoms (TaTasR) and 1.4 Å over 69 Cα atoms (SpTasH). Accessory hairpin elements are boxed and labelled: B and C in (**a**), which lack counterparts in TaTasR, and A, B, and C in (**b**), which are absent in SpTasH. The Nop and coiled-coil domains each superpose well over much of their length individually but cannot be aligned simultaneously, indicating a difference in their relative arrangement (all-pairs RMSD 4.9 Å for the Nop, 6.8 Å for the coiled-coil).

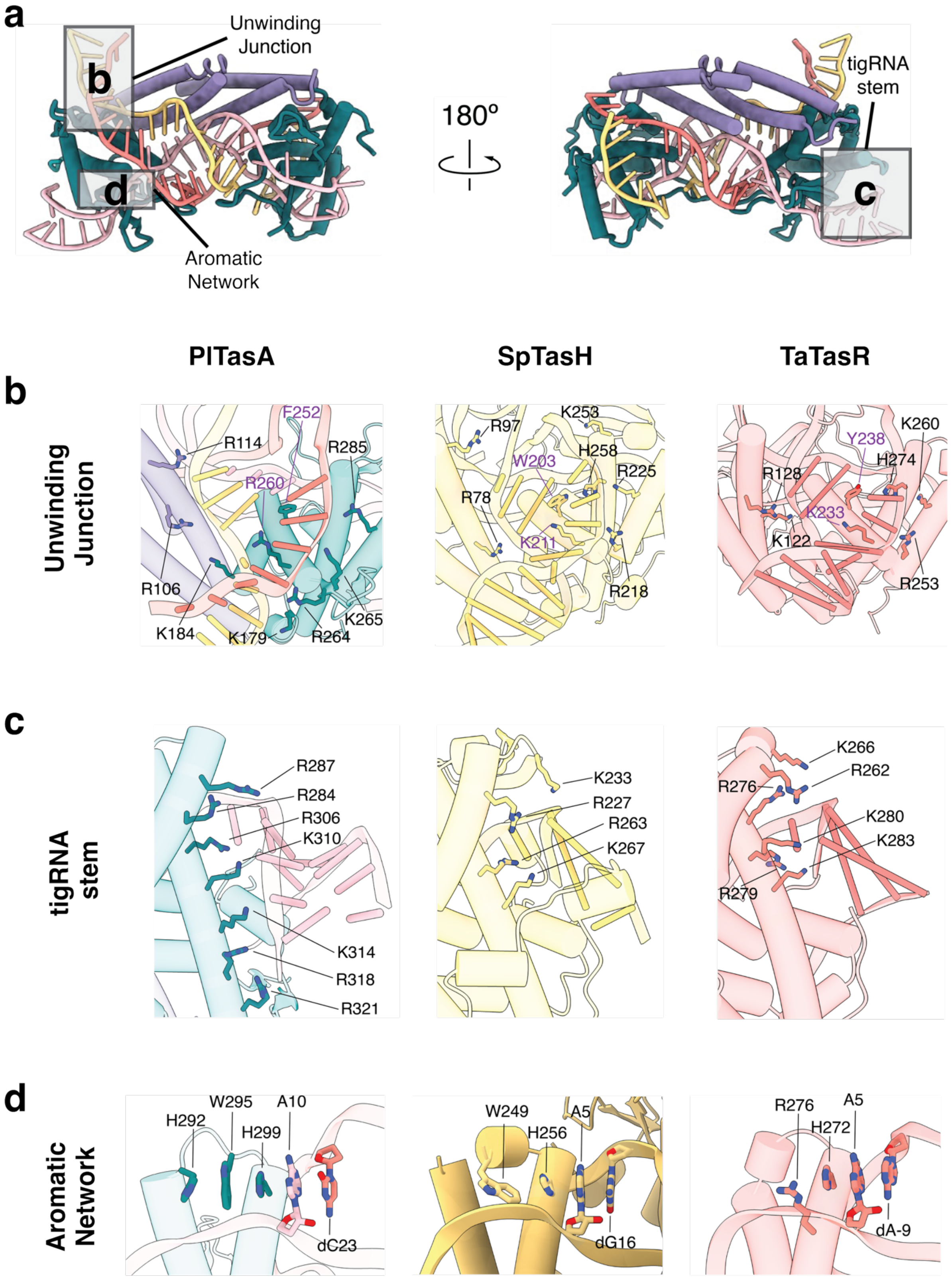
a
Unwinding Junction
b
d
Aromatic Network
180°
tigRNA stem
c
PlTasA
SpTasH
TaTasR
b
Unwinding Junction
F252
R114
R285
R260
R106
K184
K179
R264
K265
K253
R97
H258
W203
R225
R78
K211
R218
K260
Y238
H274
R128
K233
K122
R253
c
tigRNA stem
R287
R284
R306
K310
K314
R318
R321
K233
R227
R263
K267
K266
R262
R276
K280
K283
R279
d
Aromatic Network
H292
W295
H299
A10
dC23
W249
A5
H256
dG16
R276
A5
H272
dA-9

**Supplementary Figure 2. Conservation of guide and target recognition across TIGR-Tas effectors.**

**a.** PlTasA ternary complex, with the three regions detailed in b-d boxed. PlTasA is colored by domain, with the coiled-coil and Nop domains colored purple and teal, respectively.

**b.** Close-up of the unwinding junction in PlTasA (colored by domain), SpTasH (gold; PDB: 9W04), and TaTasR (salmon; PDB: 9MTY), shown left to right. An aromatic residue packs against the first paired tigRNA nucleotide and a basic residue intercalates into the duplex. The aromatic-basic pair is highlighted purple in each structure and comprises F252 and R260 in PlTasA, W203 and K211 in SpTasH, and Y238 and K233 in TaTasR.

**c.** Close-up of the tigRNA stem in the same three effectors, showing the basic residues that contact the guide backbone. In PlTasA an additional cluster (R318, R321, K314) extends along the longer stem into a region not reached in the dual-repeat systems.

**d.** Close-up of the aromatic stacking network at box C, in which a conserved adenine stacks against the DNA. The network is more extensive in PlTasA than in SpTasH or TaTasR. Key residues are shown as sticks throughout.

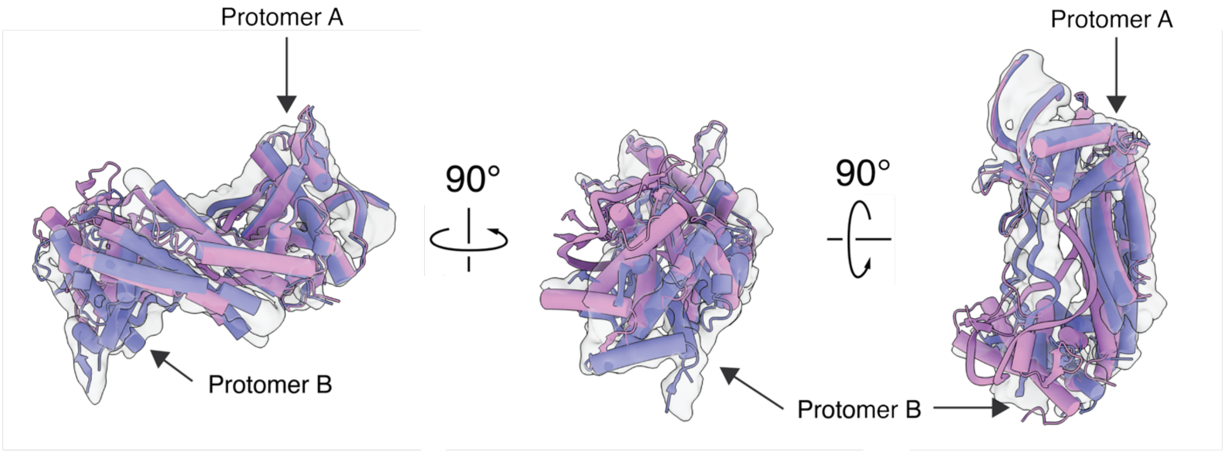


**Supplementary Figure 3. The conformational change from the binary to the ternary complex is largely restricted to protomer B.**

Superposition of the PlTasA binary (purple) and ternary (magenta) models, shown in three orientations related by 90° rotations (left to right: reference view, rotated 90° to the right, then 90° downward). The two models were superimposed over the full complex. The alignment converges on protomer A and the tigRNA stem, which remain closely superimposed between states, while protomer B undergoes a large rearrangement upon target binding. The binary complex was modeled as a backbone trace, consistent with its lower resolution.

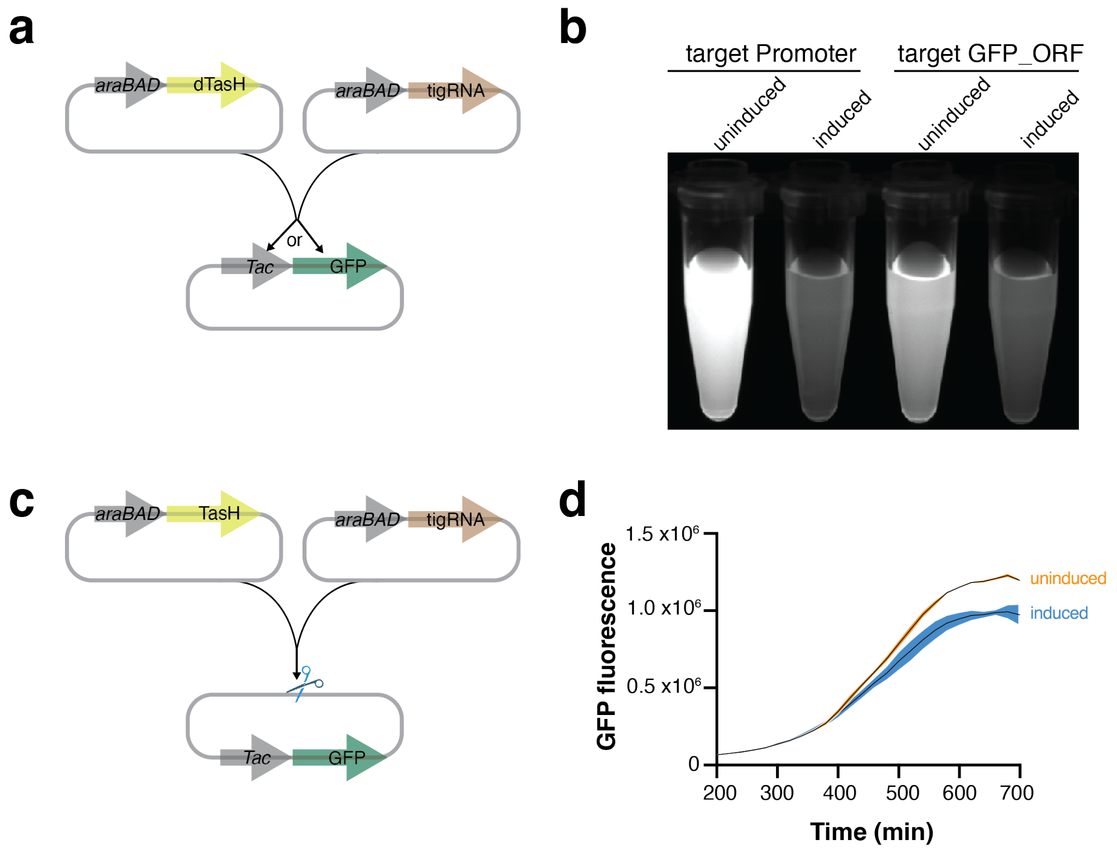


**Supplementary Figure 4. Transcriptional repression by WT and nuclease-dead SpTasH.**

**a.** Design of the GFP depletion assay. Nuclease-dead SpTasH (H54A) is targeted to the promoter or open reading frame (ORF) of a GFP reporter.

**b.** GFP fluorescence in cells expressing nuclease-dead SpTasH (H54A), showing visible reduction relative to an uninduced control.

**c.** Design of a control assay using catalytically active SpTasH.

**d.** GFP fluorescence monitored over time for uninduced and induced samples.

**a**

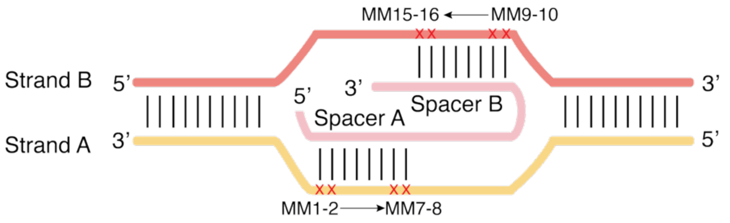


**b**

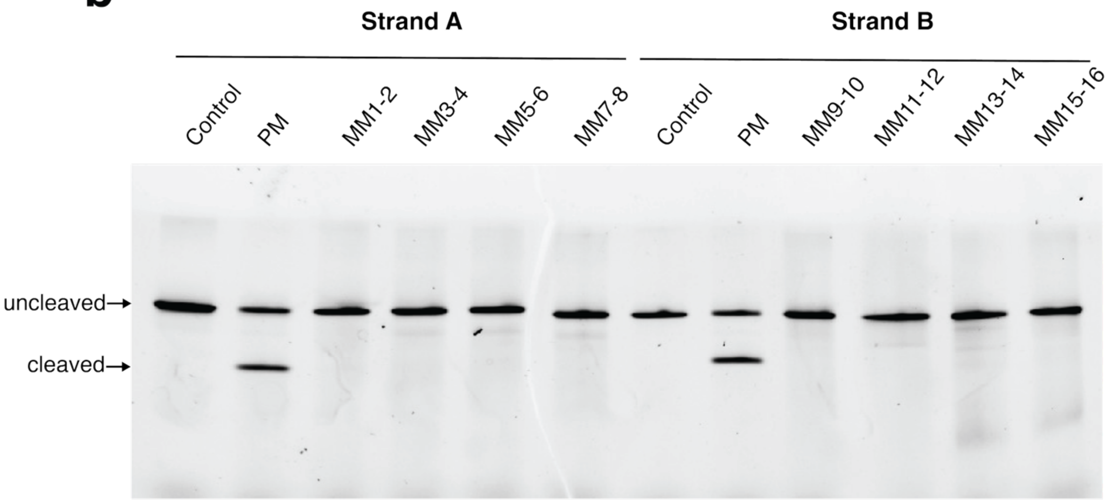


**Supplementary Figure 5. SpTasH is completely intolerant to the adjacent double mismatch.**

**a**. schematic of double mismatch design.

**b**. DNA cleavage using perfectly matched and adjacent double mismatch target. Reactions were performed for 5 hours at 37 °C.

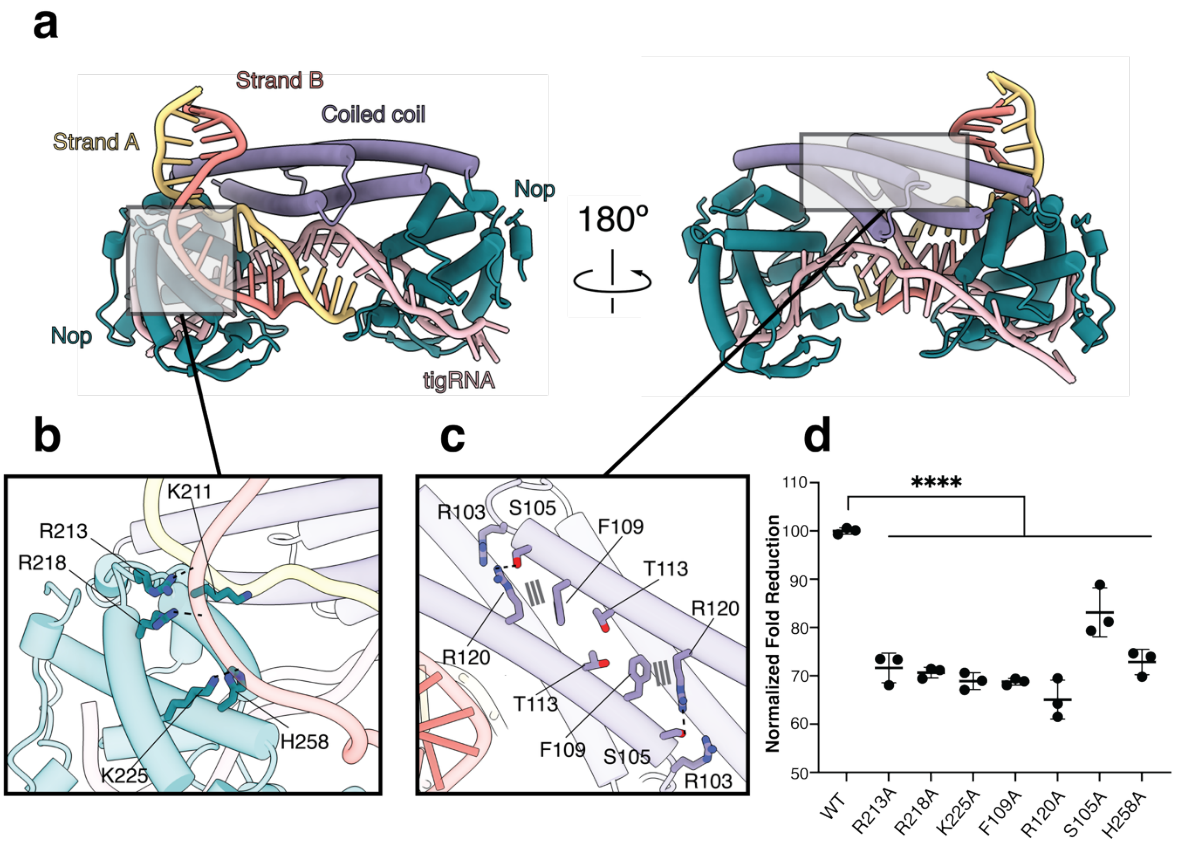


**Supplementary Figure 6. Dimerization interface and sequence-independent interactions with the DNA in SpTasH.**

**a**. Model structure of the SpTasH ternary complex at State I.

**b**. Close-up of the sequence-independent interactions with the DNA (black dashed lines denote hydrogen bonds).

**c**. Close-up of the dimerization interface between coiled coil domain (black dashed lines denote hydrogen bonds).

**d**. GFP depletion activity of SpTasH wild-type and mutants. Data represent mean $\pm$ SD (n = 3 biological replicates). Significance was determined by one-way ANOVA. ****p $\leqslant$ 0.0001.

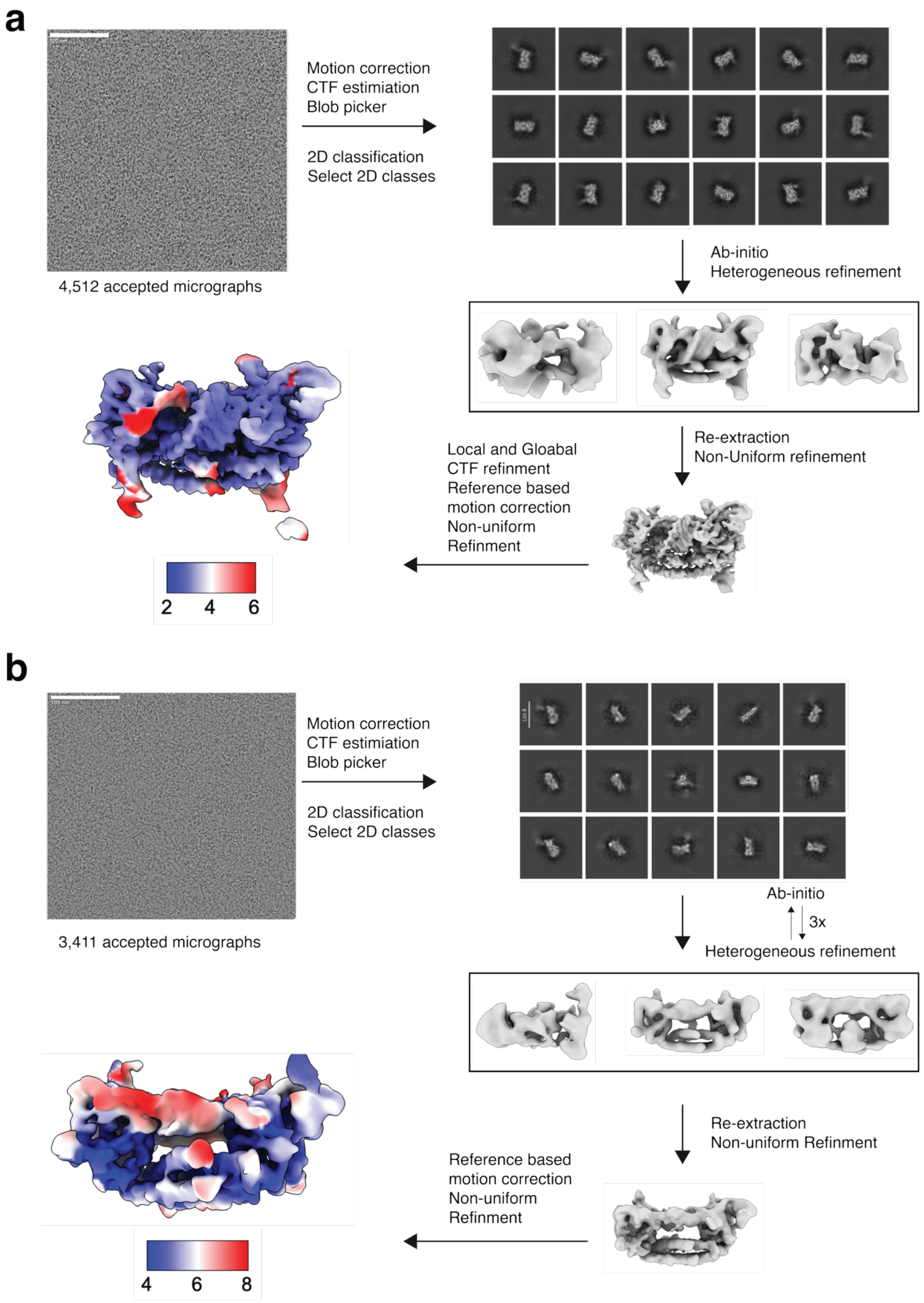


**Supplementary Figure 7. Cryo-EM data processing workflow for PITasA ternary (a) and binary (b) structures.**

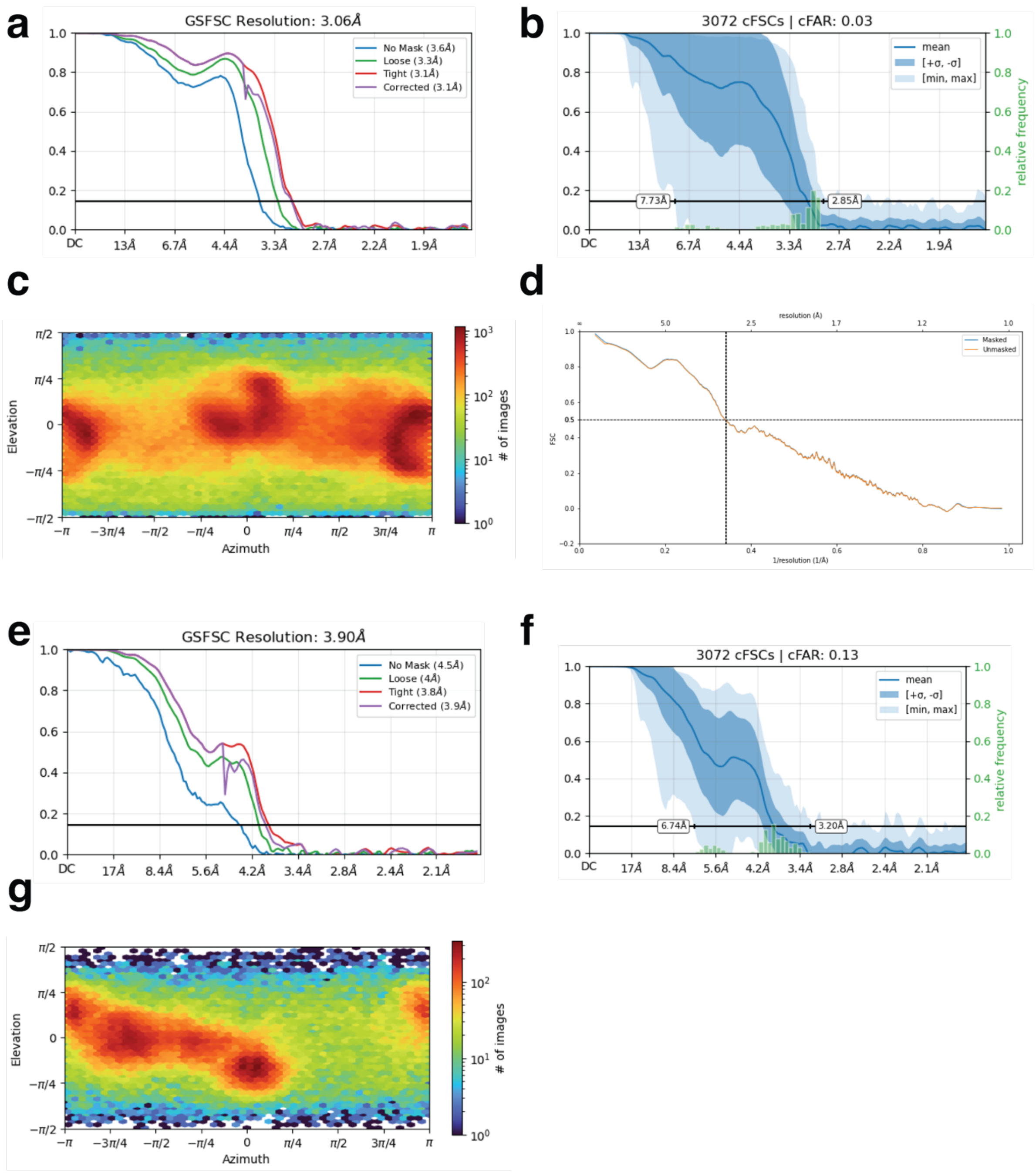


**Supplementary Figure 8. Cryo-EM data analysis of PITasA ternary (a-d) and binary (e-g) structures.**

**a**, **e**. Gold-standard Fourier Shell Correlation (FSC) curves with resolution reported at FSC = 0.143; **b**, **f**. conical FSC (cFSC) curve. **c**, **g**. Euler angle distribution plots of particle orientations; **d**. map-to-model FSC curves.

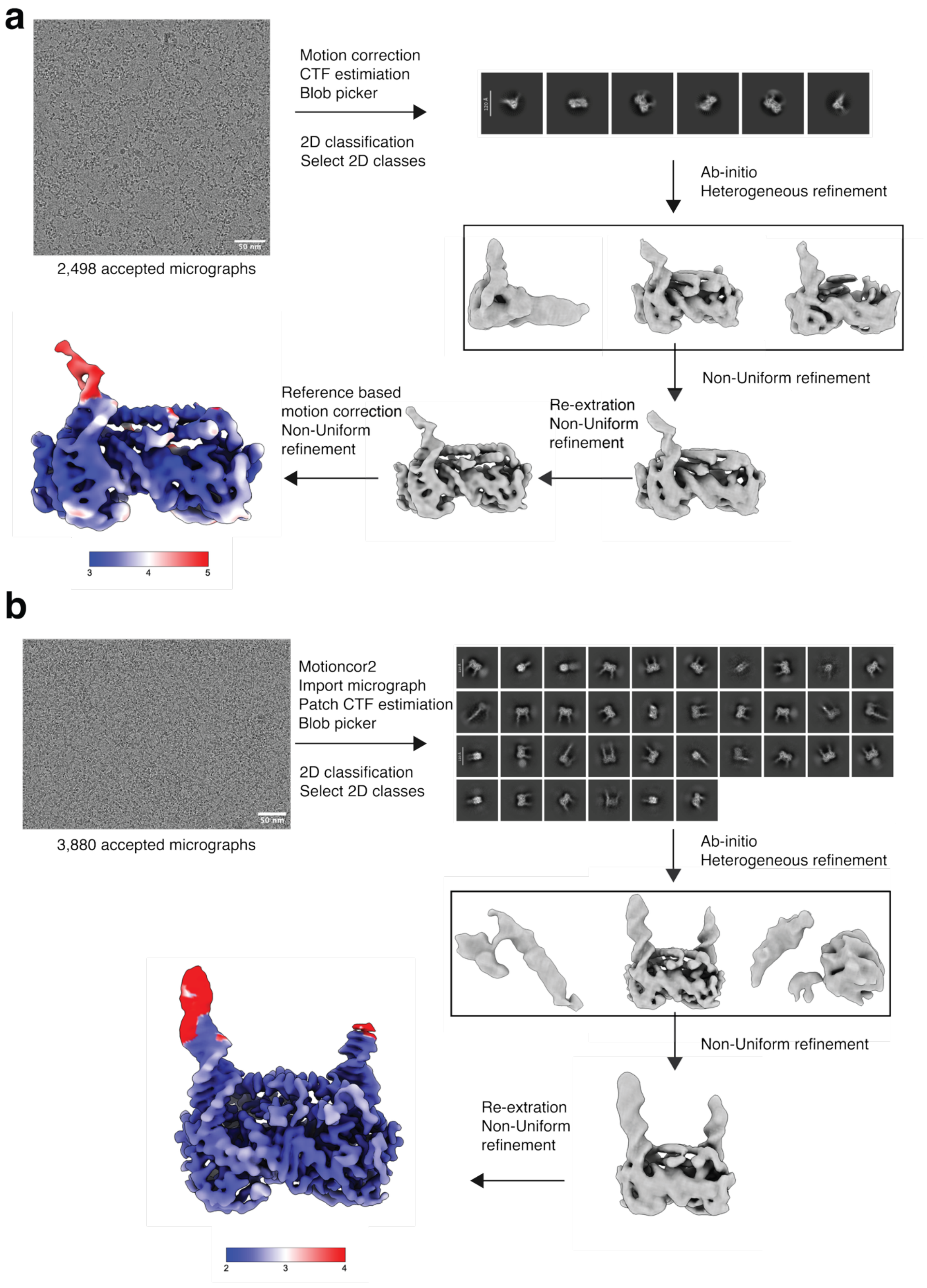


**Supplementary Figure 9. Cryo-EM data processing workflow for SpTasH State I (a) and State II (b) structures.**

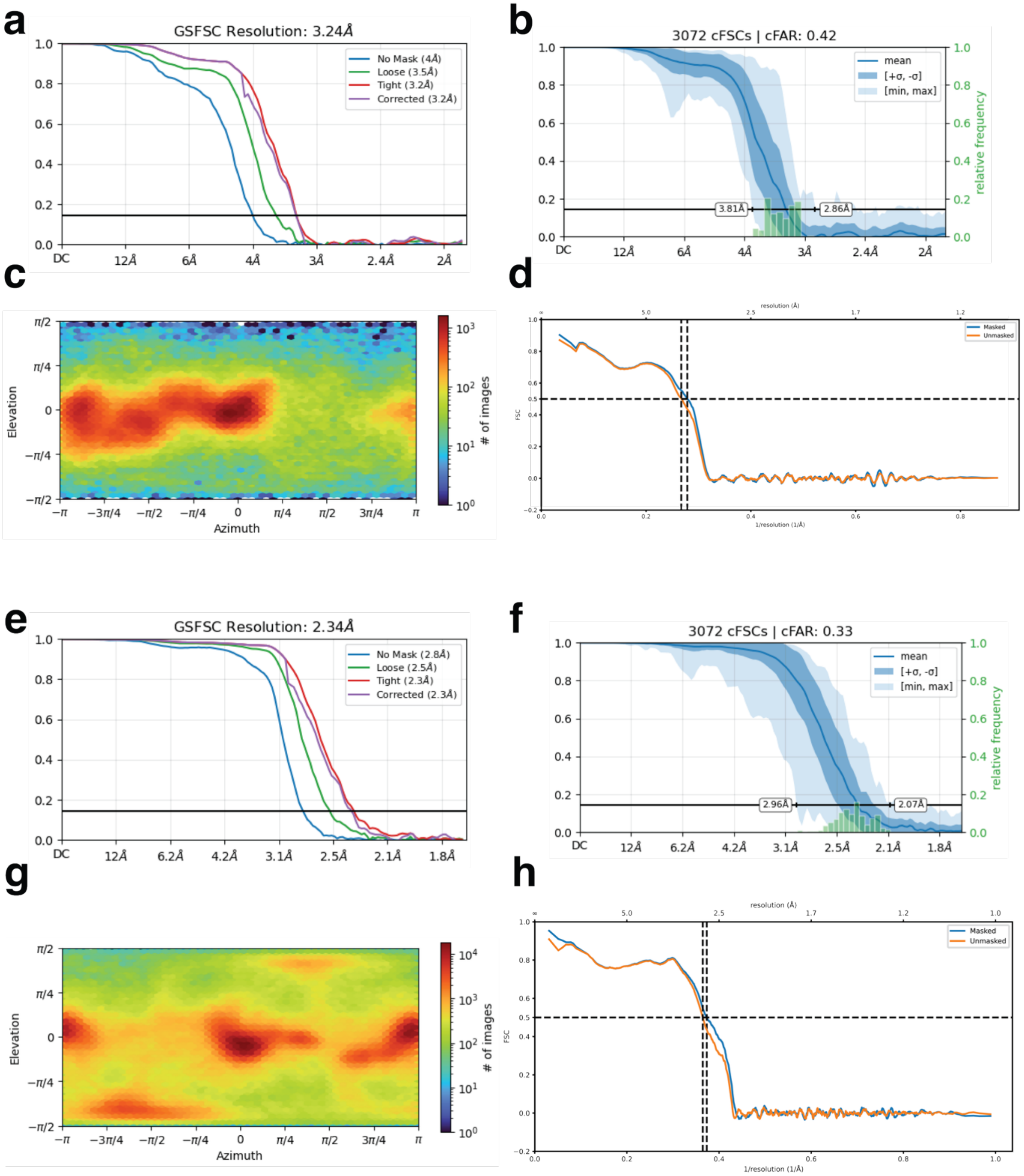


**Supplementary Figure 10. Cryo-EM data analysis of SpTasH State I (a-d) and State II (e-h) structures.**

**a**, **e**: Gold-standard Fourier Shell Correlation (FSC) curves with resolution reported at FSC = 0.143; **b**, **f**: conical FSC (cFSC) curve. **c**, **g**: Euler angle distribution plots of particle orientations; **d, h**: map-to-model FSC curves.

FpTasA-VPR

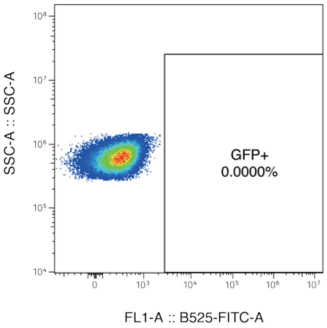


| | |
|---|---|
| Sample Name: | Untstained cells |
| Population Name: | Singlets |
| Count: | 54223 |

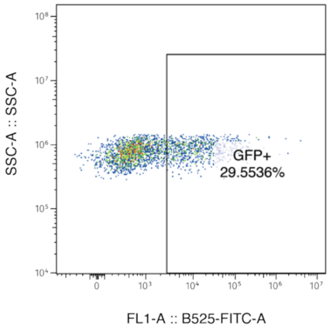


| | |
|---|---|
| Sample Name: | NT-1 |
| Population Name: | Singlets |
| Count: | 2778 |

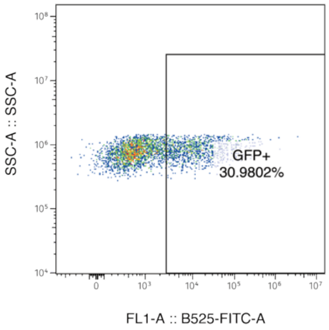


| | |
|---|---|
| Sample Name: | NT-2 |
| Population Name: | Singlets |
| Count: | 3428 |

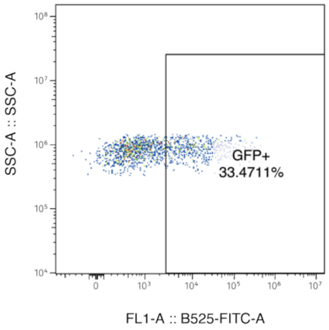


| | |
|---|---|
| Sample Name: | NT 3 |
| Population Name: | Singlets |
| Count: | 1452 |

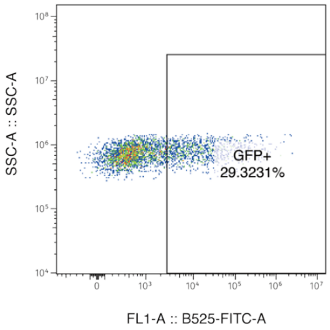


| | |
|---|---|
| Sample Name: | SV40-T1-1 |
| Population Name: | Singlets |
| Count: | 3250 |

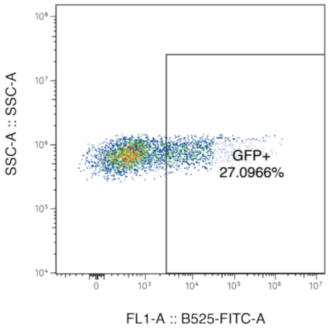


| | |
|---|---|
| Sample Name: | SV40-T1-2 |
| Population Name: | Singlets |
| Count: | 3768 |

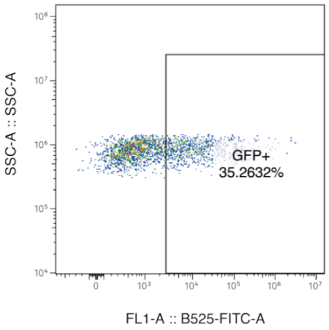


| | |
|---|---|
| Sample Name: | SV40-T1-3 |
| Population Name: | Singlets |
| Count: | 2280 |

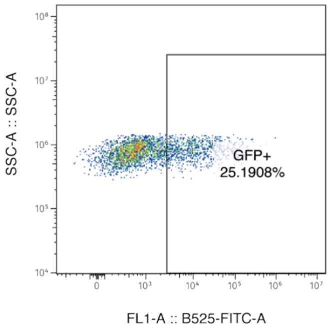


| | |
|---|---|
| Sample Name: | SV40-T2-1 |
| Population Name: | Singlets |
| Count: | 3537 |

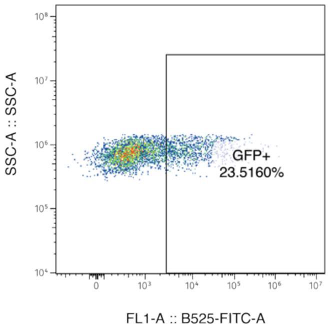


| | |
|---|---|
| Sample Name: | SV40-T2-2 |
| Population Name: | Singlets |
| Count: | 3959 |

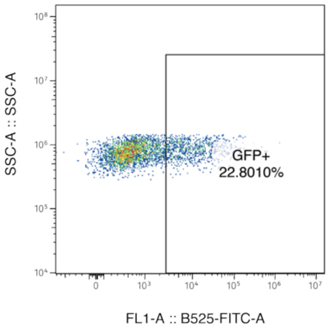


| | |
|---|---|
| Sample Name: | SV40-T2-3 |
| Population Name: | Singlets |
| Count: | 3513 |

**Supplemental Figure 11: Flow cytometry analysis of GFP-positive HEK293T cells expressing FpTasA-VPR.**

Untransfected cells were used to establish the gating threshold for GFP-positive cells. The percentage of GFP-positive cells was determined for cells co-transfected with FpTasA-VPR and NT, SV40-T1, or SV40-T2 tigRNA constructs. A total of 10,000 events were recorded for each sample.

PlTasA-VPR

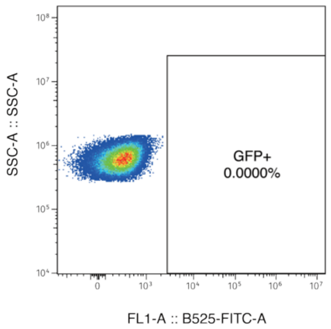


Sample Name: Untstained cells
Population Name: Singlets
Count: 54223

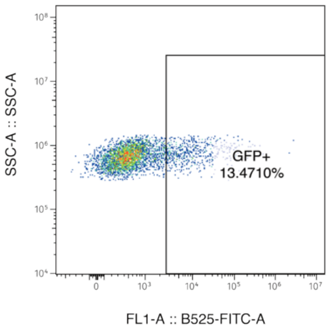


Sample Name: NT-1
Population Name: Singlets
Count: 3875

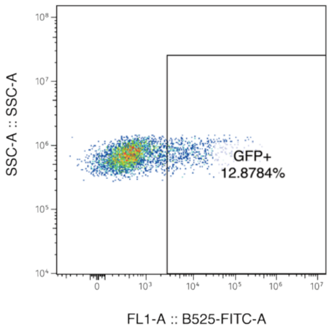


Sample Name: NT-2
Population Name: Singlets
Count: 4162

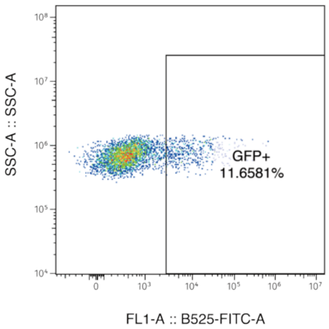


Sample Name: NT-3
Population Name: Singlets
Count: 4083

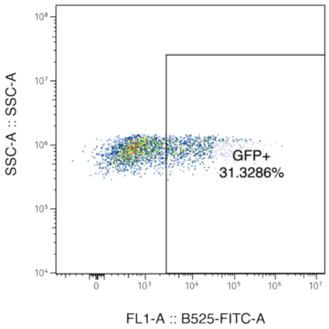


Sample Name: SV40-T1-1
Population Name: Singlets
Count: 2544

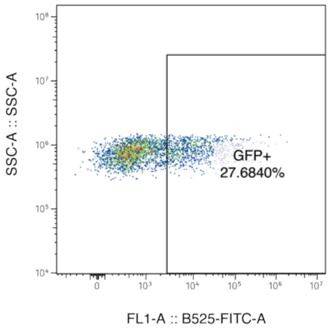


Sample Name: SV40-T1-2
Population Name: Singlets
Count: 3139

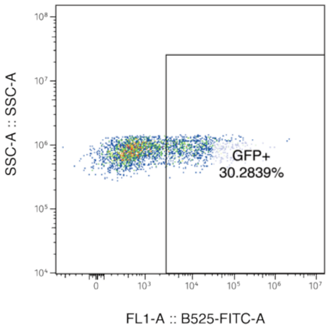


Sample Name: SV40-T1-3
Population Name: Singlets
Count: 2853

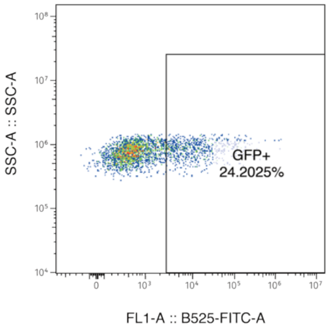


Sample Name: SV40-T2-1
Population Name: Singlets
Count: 2884

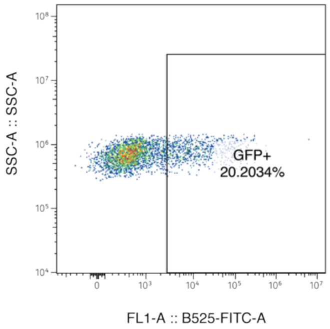


Sample Name: SV40-T2-2
Population Name: Singlets
Count: 3737

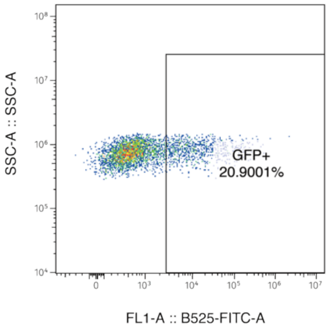


Sample Name: SV40-T2-3
Population Name: Singlets
Count: 3555

**Supplemental Figure 12: Flow cytometry analysis of GFP-positive HEK293T cells expressing PITasA-VPR.**

Untransfected cells were used to establish the gating threshold for GFP-positive cells. The percentage of GFP-positive cells was determined for cells co-transfected with PITasA-VPR and NT, SV40-T1, or SV40-T2 tigRNA constructs. A total of 10,000 events were recorded for each sample.
